# Future shapes present: autonomous goal-directed and sensory-focused mode switching in a Bayesian allostatic network model

**DOI:** 10.1101/2024.04.03.588025

**Authors:** Hayato Idei, Jun Tani, Tetsuya Ogata, Yuichi Yamashita

**Affiliations:** Department of Information Medicine, National Institute of Neuroscience, National Center of Neurology and Psychiatry, Tokyo, 187-8502, Japan; Cognitive Neurorobotics Research Unit, Okinawa, Okinawa Institute of Science and Technology, 904-0495, Japan; Department of Intermedia Art and Science, Waseda University, Tokyo, 169-8555, Japan

**Keywords:** Autonomy, Mode switching, Intention, Allostasis, Self-organization, Entropy, Variational Bayes, Free-energy principle

## Abstract

Trade-offs between moving to achieve goals and perceiving the surrounding environment highlight the complexity of continually adapting behaviors. The need to switch between goal-directed and sensory-focused modes, along with the goal emergence phenomenon, challenges conventional optimization frameworks, necessitating heuristic solutions. In this study, we propose a Bayesian recurrent neural network framework for homeostatic behavior adaptation via hierarchical multimodal integration. In it, the meta-goal of “minimizing predicted future sensory entropy” underpins the dynamic self-organization of future sensorimotor goals and their precision regarding the increasing sensory uncertainty due to unusual physiological conditions. We demonstrated that after learning a hierarchical predictive model of a dynamic environment through random exploration, our Bayesian agent autonomously switched self-organized behavior between goal-directed feeding and sensory-focused resting. It increased feeding before anticipated food shortages, explaining predictive energy regulation (allostasis) in animals. Our modeling framework opens new avenues for studying brain information processing and anchoring continual behavioral adaptations.

## 1. Introduction

Continual behavior adaptation in an ever-changing world involves continuous mode switching between moving to achieve a goal (“goal-directed”) and sensitively perceiving their environment (“sensory-focused”). Locomotion for obtaining food or water and examining the surrounding environment or danger clearly demonstrates the significance of the dual modes for survival. Notably, cognitive agents exhibit predictive regulation of nutritional (energy) status in anticipation of future environmental changes^1–3^, suggesting that the predicted future shapes present behavior. For example, hibernating or migrating animals increase their food intake prior to periods when food is scarce or unavailable^4,5^. This phenomenon, referred to as behavioral allostasis, shows that both modes play a profound role in maintaining homeostasis in a dynamic environment. However, trade-offs between goal-directed locomotion and sensory-focused perception (e.g., to carefully hear the surrounding sounds, one must stop) as well as the phenomenon of goal emergence complicate the mechanistic understanding of autonomous mode switching. How does the brain resolve these challenges and autonomously adapt to behavior^6,7^?

Autonomous mode switching between both modes exposes critical issues in the computational modeling of the brain and the construction of autonomous intelligence. The notion that the brain is a predictive or generative machine has been formalized in neuroscience as predictive coding and active inference (or the free-energy principle)^8–12^. Additionally, it is applied to architectures in machine learning (deep learning) and robotics^13–15^ as well as computational understanding of psychiatric disorders^16–20^. The neurobiologically and systematically plausible computational basis converges into the Bayesian framework, or prediction error minimization (minimization of the mismatch between real and predicted sensations). Within which a Bayesian agent infers the causes of sensations as internal beliefs (latent variables), influenced by prior experiences. The conflict between the computational processes of perception and movement is a critical issue in explaining autonomous switching between both modes. Specifically, in the context of perception (predictive coding), Bayesian beliefs should be adapted to reduce sensory prediction errors by fitting sensations. In contrast, during the execution of movements (active inference), these beliefs should remain unchanged and be realized by changing sensations. In other words, Bayesian beliefs serve as “recognition” in sensory-focused perception but “intention” in goal-directed movement. Furthermore, goal-directed movement itself is a risk factor for prediction errors due to interactions with the environment. This may be related with the argument of “the dark-room problem”^21^.

Previous computational modeling and neurorobotic studies have heuristically avoided these problems by operating or modeling one process of movement or perception at a time^22,23^, explicitly training an agent to move in a specific manner^24–26^, and/or preparing separate latent variables and cost functions for perception and movement (action planning)^27–29^. However, these heuristic solutions neglect autonomous mode switching between goal-directed movement and sensory-focused perception in the face of computational conflicts experienced in daily life. Furthermore, the same latent variable (Bayesian belief) can serve as both “recognition” and “intention” (e.g., the experience of seeing or recognizing a certain food in a forest leads to the formation of the intention to find that food in the forest later on). Behavioral allostasis through autonomous switching between both modes may require an account beyond prediction error minimization for explaining how “recognition” changes into “intention” and vice versa. This may involve the dynamic self-organization of future sensorimotor goals and their precision (or strength), which modulates the balance between sensory information and goal representation.

In this study, we propose that a cognitive agent not only minimizes the sensory prediction errors at hand but also minimizes the predicted future sensory entropy (uncertainty). This hypothesis aims to explain the continual adaptation of homeostatic behaviors by linking the Bayesian brain hypothesis with a physical view of homeostasis. As physical systems, biological cognitive agents naturally decay without a cycle of nutrition or excretion, eventually leading to death. This inevitable tendency toward disorder may be associated with an increase in entropy owing to irreversible physical processes within the system (body)^30^. Thus, homeostatic behaviors of cognitive agents may be regarded as means of resisting the tendency toward disorder, as Schrödinger originally stated that life feeds on “negative entropy” or free-energy^31^. We consider this physical constraint in brain information processing given that sensation is the interface between the brain information processing system and the bodily physical system. Then, minimization of predicted future sensory entropy can be considered a “meta-goal” concerning second-order prediction (of uncertainty), which may explain the dynamic self-organization of future sensorimotor goals and their precision or strength for behavioral allostasis.

The basic premise of this hypothesis is that appropriate physiological conditions in the body are required to maintain homeostasis. Conversely, unusual physiological conditions increase physiological entropy. This may be supported by that certain food habits (e.g., inappropriate caloric content of diet) and thermal stress increase entropy production^32–35^. Sensory (information) entropy, or uncertainty, is a potential source from which the brain can recognize the homeostatic context of the system.

To test our hypothesis, we developed a Bayesian neural network framework for homeostatic behavioral adaptation via hierarchical multimodal integration. In addition, we showed that the meta-goal of Bayesian allostasis (minimizing entropy) can be derived by extending the framework of variational Bayes, a formal tractable solution of Bayesian inference, into future predictive processing. Our simulation experiment demonstrates how autonomous mode switching and recognition-intention transitions emerge through Bayesian allostasis. Our modeling framework provides a basis for exploring continual behavioral adaptation and the underlying computational mechanisms in the brain network.

## 2. Results

### 2.1. Constraints on environment and sensation

We designed a navigation survival task in a dynamic environment in which a cognitive agent was required to survive through predictive interoceptive (energy) regulation. In the environment of the simulated square space [−1.0,1.0] × [−1.0,1.0], food was placed at a random position (Fig. 1a). The agent’s energy state increased, if the agent approached it. In a 100 timestep cycle, the nutritional value of the food gradually decreased, and the food reappeared at another random position (Fig. 1b). It roughly mimics the changes in locations and the amount of food in the natural environment. In addition, a cue was placed at a fixed position (0.0, 0.4) that temporarily informed the agent of the x-y coordinate of the current food position if the agent approached it. This cue setting was inspired by a previous modeling study^36^.

**Fig. 1:**
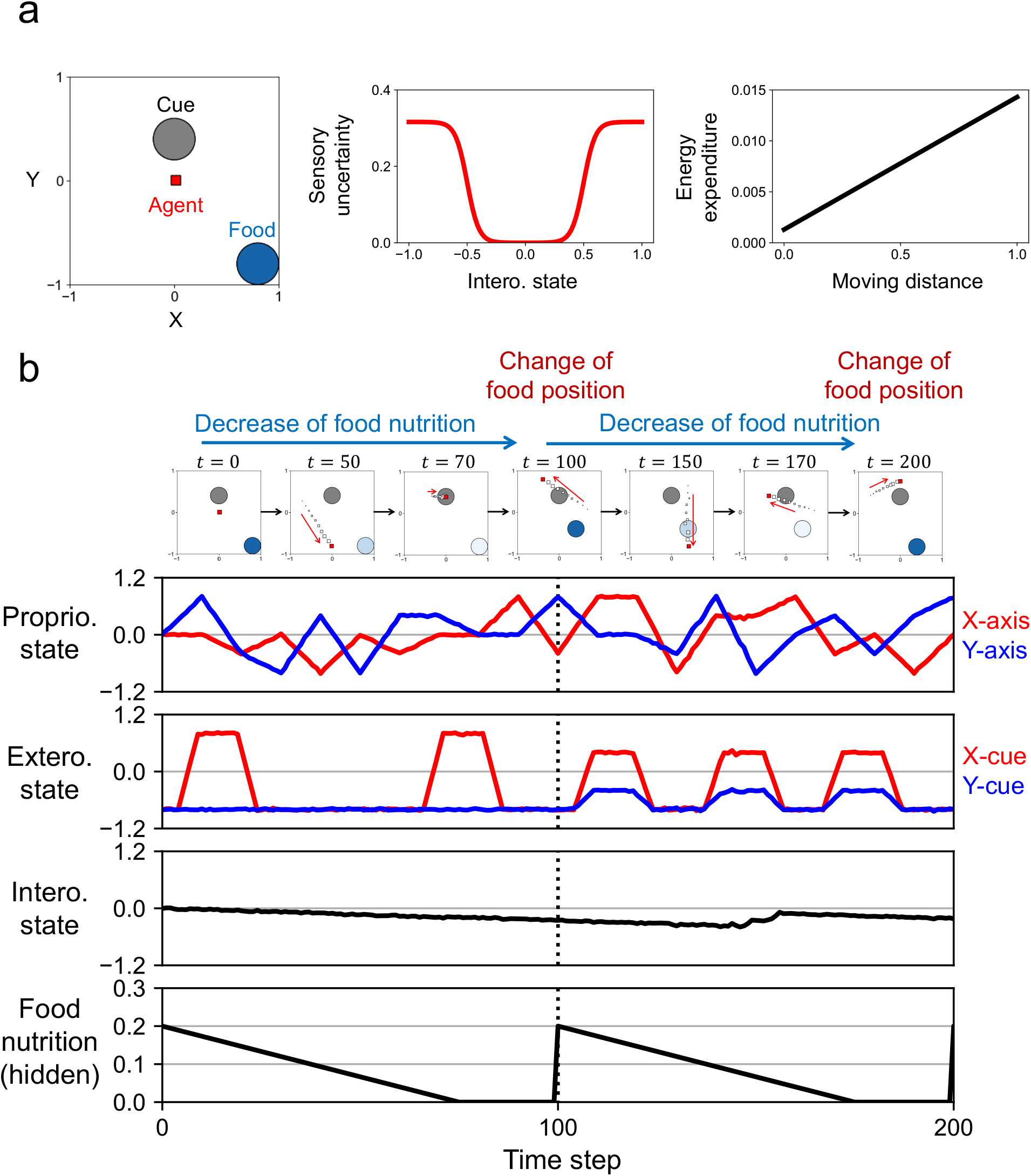
Constraints on environment and sensation. **(a)** Environmental setup. In the environment of simulated square space [−1.0,1.0] × [−1.0,1.0], a food is placed at a random position, and a cue is placed at a fixed position (0.0, 0.4). Unusual interoceptive state causes large sensory (information) uncertainty, and movements lead to energy expenditure. **(b)** Timeseries of proprioceptive state, exteroceptive state, interoceptive state, and food nutrition during random-exploration learning. Note that the agent does not sense the current food position or food nutrition directly. Each sensation takes values between -1.0 and 1.0, corresponding to the range of neural network output. The first 200 steps of learning data (100,000 steps) are shown.

At each time step, the agent received two-dimensional x-y coordinates of the body position, a two-dimensional cue signal, and a one-dimensional energy state, which were regarded as proprioceptive, exteroceptive, and interoceptive sensations, respectively. Each sensation took a value between -1.0 and 1.0, corresponding to the range of the neural network output. The exteroceptive state is meaningful only when the agent is near the cue, taking the value (−0.8, -0.8) when it is not. If the food is located at (−0.8, -0.8), the agent can deduce this because the exteroceptive state remains unchanged while near the cue. As a physiological constraint of the agent, a low or high interoceptive state (negative or positive deviations from 0.0 value) causes large sensory (information) uncertainty implemented as Gaussian noise added to all sensory modalities (Fig. 1a). We assume that this corresponds to the effects of deviations from homeostasis or an increase in physiological entropy due to excessively low or high interoceptive states. We regard the agent as dead when the interoceptive state becomes -1.0 or 1.0. In addition, we assume that the movements of the agent lead to energy expenditure (a decrease in the interoceptive state).

#### Multimodal Bayesian homeostatic recurrent neural network

For behavioral allostasis, the agent must have an internal predictive model of the environment that represents how multimodal sensations and their uncertainties are changed by internal and external causes. Here, we propose the multimodal Bayesian homeostatic recurrent neural network (MBH-RNN), which integrates multimodal sensorimotor processing and homeostatic processing based on variational Bayes (Fig. 2a). The MBH-RNN has a structural and temporal hierarchy consisting of sensorimotor modules (lower-perceptual modules distributed for each modality and multimodal-associative modules) and homeostatic modules (unexpected-uncertainty cause module and higher-cognitive module). Sensorimotor modules process information from each sensory modality and its associations. They may be associated with the functionality of brain regions such as sensory areas and the insula cortex^37^. In contrast, the higher-cognitive module integrates multimodal information with uncertainty (homeostatic) information, which may be analogous to cognitive control regions in the brain, such as the prefrontal and anterior cingulate cortices, which are thought to control a wide range of motivational, goal-directed, and uncertainty-related behaviors^38,39^. Moreover, the unexpected-uncertainty-cause module is primarily involved in the perception of the cause of uncertainty, including unexpected causes, and generates predictions about sensory uncertainty related to all sensory modalities. The function of this module is essential for computing the predicted future sensory entropy related with the homeostatic context and driving feeding behavior through interactions with the higher-cognitive module. We hypothesize that the unexpected-uncertainty-cause module may be associated with brain regions, such as the amygdala and basal forebrain, which are thought to represent a range of uncertainty information, including unexpected uncertainty, and modulate feeding behavior^40–44^.

**Fig. 2:**
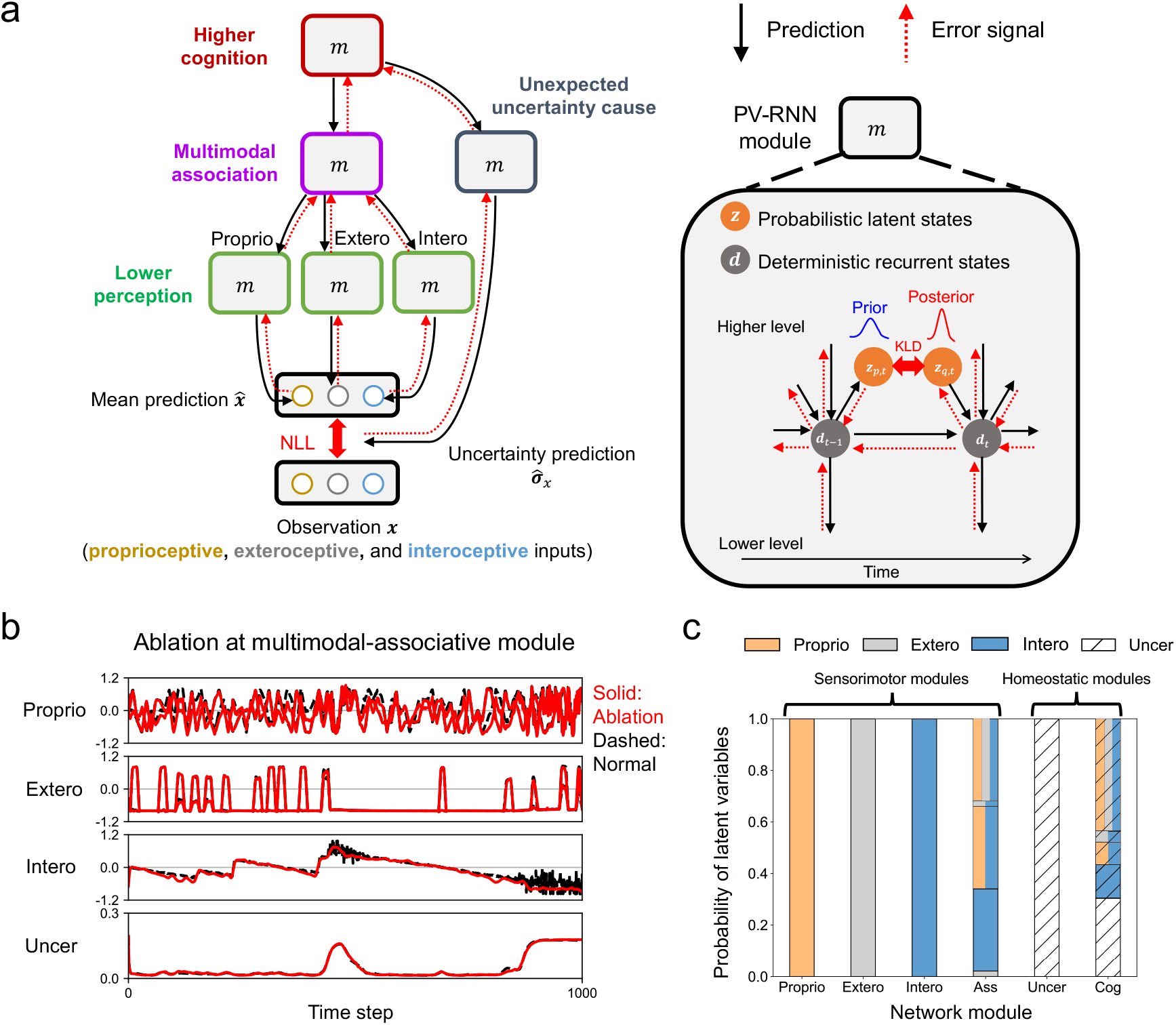
Multimodal Bayesian homeostatic recurrent neural network. Structure and information processing of multimodal Bayesian homeostatic recurrent neural network (MBH-RNN). Each module *m* is based on a predictive-coding-inspired variational RNN (PV-RNN). Prior and posterior distributions of latent state are represented as single Gaussian distributions for clarity, although each of them is a multivariate Gaussian distribution. NLL: negative log-likelihood. KLD: Kullback-Leibler divergence. **(b)** An example of the effectof ablating a latent variable in the multimodal-associative module on generations of predictions about proprioceptive state, exteroceptive state, interoceptive state, and sensory uncertainty. **(c)** Probability of trained latent variables representing either proprioceptive, exteroceptive, interoceptive, or sensory uncertainty information. A simultaneous display of colors or lines means the representation of multiple pieces of information. Proprio: proprioceptive module. Extero: exteroceptive module. Intero: interoceptive module. Ass: multimodal-associative module. Uncer: unexpected-uncertainty-cause module. Cog: higher-cognitive module.

Each module of the MBH-RNN is based on a predictive-coding-inspired variational RNN (PV-RNN)^45,46^. A brief explanation of the information processing in the MBH-RNN is as below (see Fig. S1 and the Methods section for details). At each time step *t*, the latent states 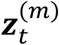 of the MBH-RNN represent Bayesian beliefs regarding the causes of sensations or their uncertainty as (multivariate) Gaussian distributions individually assigned to each module *m*. Based on the Bayesian brain hypothesis^8^, each latent state has prior and posterior probability distributions that correspond to the estimated hidden causes before and after observing the current sensations, respectively. Based on the latent states, the MBH-RNN generates top-down predictions about the mean 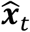 and standard deviation (uncertainty) 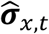 of sensations ***x***_*t*_ as a Gaussian distribution. Here, the deterministic recurrent states 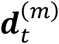 transform latent states into sensory predictions via synaptic connections that represent the dynamic relationships between sensations and their causes. The MBH-RNN uses a multiple timescale RNN (MT-RNN) as the transformation function, which represents the temporal hierarchy by a multiple timescale property in neural activation, inspired by the biological brain^24^. Specifically, higher-level modules (e.g., multimodal-associative module) have slower neural dynamics than lower-level modules have (e.g., lower-perceptual modules). Importantly, 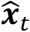 and 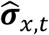 are generated through different network pathways, where the higher-cognitive module serves as the information hub. This distinction of the prediction pathways enables us to dissociate the sensorimotor processing from homeostatic processing and assures that the predicted sensory uncertainty 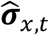 is the uncertainty caused by homeostatic context that affects all sensory modalities.

The cost function of the MBH-RNN is variational free-energy (VFE) that is introduced as a tractable quantity within variational Bayes that bounds the surprisal (or the negative log model evidence) for sensations^46^ (see Methods for the derivation).

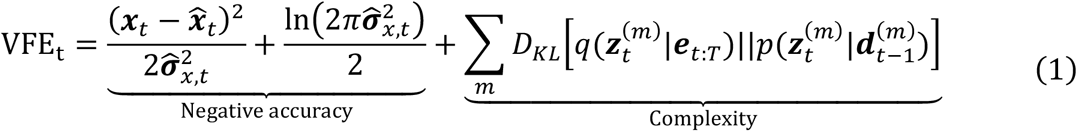

The first term (negative accuracy term) of the VFE describes the negative log-likelihood (NLL) or precision-weighted sensory prediction error. The second term (complexity term) describes Kullback-Leibler divergence (KLD) between the posteriors 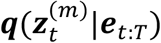 and priors 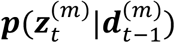. The priors are generated from prior experience through previous deterministic recurrent states 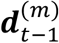 whereas the posteriors are determined by the backpropagated errors ***e***_*t*:*T*_ (*T*: the last time step of the time window).

#### Learning of internal predictive model

The learning of the MBH-RNN was performed using data acquired through random exploration of the environment for 100 food nutrition cycles (10,000 time steps). Fig. 1b shows the time series of the proprioceptive, exteroceptive, and interoceptive sensations during random exploration. The learning data did not contain any organized movements, and the proprioception data for random movements were generated automatically using a predefined algorithm. The MBH-RNN learned to reconstruct the experienced sensations ***x***_1:*T*_ (*T*=10,000) by repeating the prediction generations and parameter updates within time steps 1: *T* 100,000 times offline (Fig. S1). In the learning process, posteriors at each time step ***z***_***q***,1:*T*_ and synaptic weights ***ω*** are updated through the gradient descent method for the accumulated VFE over time steps, where the partial derivative of the VFE with respect to each parameter is calculated by the back propagation through time (BPTT) algorithm^47^.

We investigated how latent states in the trained MBH-RNN represented multimodal information. It was analyzed through an ablation study by removing each latent variable one-by-one and evaluating the influence on the generations of 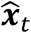 and 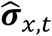 (see Methods for the evaluation). Fig. 2b shows an example of the effect of ablating a latent variable in the multimodal-associative module. The ablation of the latent variable impacted the generation of predictions about the proprioceptive and interoceptive states (the solid red lines deviate from the black dashed lines), but not those about the exteroceptive state and sensory uncertainty. This suggests that the ablated latent variable represents bimodal proprioceptive and interoceptive information. Based on the ablation analysis for all latent variables, we categorized the latent variables depending on the information they represented. As shown in Fig. 2c, each of the lower-perceptual modules developed a corresponding modality-specific representation, whereas the multimodal-associative module developed bimodal and trimodal latent variables and represented interoceptive information and its integration with proprioceptive and exteroceptive information. However, the unexpected-uncertainty-cause module was only involved with sensory uncertainty predictions, while the higher-cognitive module represented the integration of multimodal information and sensory uncertainty. This suggests that the unexpected-uncertainty-cause module represents the estimated cause of sensory uncertainty that is independent of the internal and external environmental states, whereas the higher-cognitive module represents the internal and external environmental causes of sensory uncertainty. Thus, we confirmed that the MBH-RNN can be used to develop a hierarchical multimodal predictive model of the environment.

#### Autonomous goal-directed and sensory-focused mode switching

Using the trained MBH-RNNs, we analyzed the autonomous behavior generation of the agent in a dynamic environment, in which the random changes in food position in a 100 timestep cycle were different from the learning experience. Ten test trials were performed by each of the 10 trained MBH-RNNs. Autonomous behavior generation was implemented as a simultaneous operation of perception, action generation, and goal modulation based on VFE minimization from past to future (Fig. 3a and S2). Specifically, at the current sensorimotor time step *t*_*c*_ (≥ 0), the MBH-RNN, with fixed synaptic weights, repeats prediction generations and posterior updates within time steps from *t*_*c*_ − *win*_*f*_ + 1 to *t*_*c*_ + *win*_*f*_ for 200 times in an online manner. The length of the past time window is *win*_*p*_ = 10 (or *win*_*p*_ = *t*_*c*_ + 1 *if t*_*c*_ < 10<, while that of the future time window is *win*_*f*_ = 200. The cost function in the past time window is the accumulated VFE of the past, which is calculated based on the observed sensations 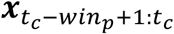.

**Fig. 3:**
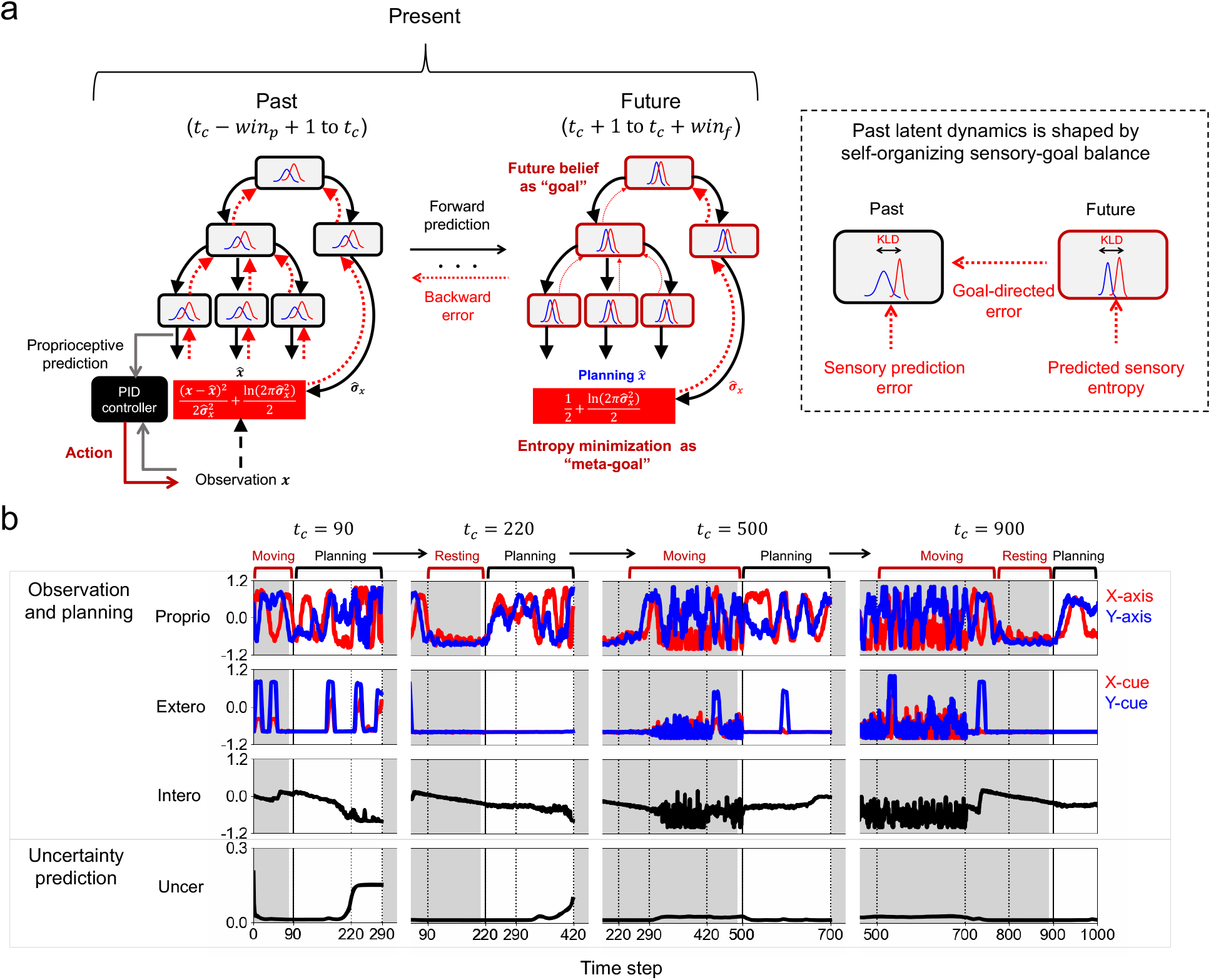
Autonomous Bayesian allostasis model. **(a)**Online inference of latent states is performed via minimization of variational free energy from the past to the future. *t*_*t*_, *win*_p_, and *win*_*f*_ indicate the current sensorimotor time step, the length of past time window, and the length of the future time window, respectively (*win*_p_ = 10, *win*_*f*_ = 200). Action is generated via a proportional-integral-derivative (PID) controller based on proprioceptive prediction at the current sensorimotor time step *t*_*c*_. Latent states in the past were determined through interaction among previous experiences, sensory prediction errors, and goal-directed errors. KLD: Kullback-Leibler divergence. **(b)** An example of timeseries for proprioceptive state (proprio), exteroceptive state (extero), interoceptive state (intero), and predicted sensory uncertainty (uncer) during an autonomous survival task. The estimated sensory sigma is the average value over all sensations. The white-colored background indicates the time window of network predictive processing from the past to the future. In the future time window, the proprioceptive, exteroceptive, and interoceptive states are predicted values (planning), not real sensations. See Fig. S3 for the corresponding timeseries of latent states.

This drives the postdictive perceptual inference of internal and external states. In addition, for future predictive processing, we extend the VFE into the future time window while considering that sensations in the future are unknown variables. Inspired by a previous mathematical study^48^, we derive a form of the variational free energy of the future (VFEF) by evaluating variational free energy under the expectation of predictions about future sensations (that bounds the expected surprisal, or the expected negative log model evidence). This changes the NLL term of the VFE into the predicted conditional (Shannon) entropy of sensation (see Methods for derivation).

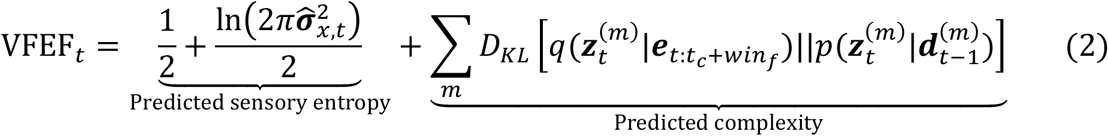

Here, 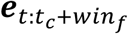 indicates the back-propagated errors calculated using the BPTT algorithm, which modulates the posteriors (*t*_*c*_ + *win*_*f*_ is the last time step of the future time window). This derivation of the VFEF naturally describes the meta-goal of the proposed Bayesian allostasis (the minimization of entropy). At each sensorimotor time step *t*_*c*_, the MBH-RNN updates the posteriors 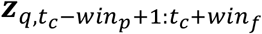 to minimize the accumulated free-energy *F*_*allostasis*_ through BPTT.

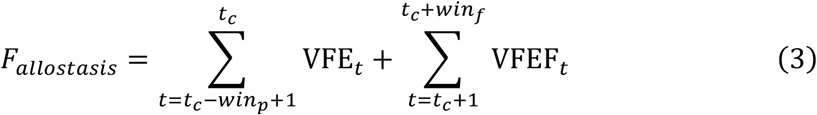

An important aspect of the posterior updates through the back propagation is that the past posteriors are determined through interactions among previous experiences, sensory prediction errors, and goal-directed errors (Fig. 3a), in which we regard future beliefs as future “goal” that is adapted towards minimizing predicted future sensory entropy. The action generation of the agent is performed via the realization of proprioceptive predictions generated at the current sensorimotor time step *t*_*c*_ before the posterior updates using a proportional-integral-derivative (PID) controller. We regard sequential future predictions about proprioceptive, exteroceptive, and interoceptive sensations as “planning.”

Fig. 3b shows an example of how the agent achieved continual behavioral adaptation for homeostasis in the test trial (see also Fig. S3 and Video S1). At *t*_*c*_ = 90 in Fig. 3b, the agent kept the interoceptive state at approximately 0.0 (appropriate level) by previously moving for food intake. The MBH-RNN planned to continue moving (see predicted proprioceptive states in future planning). In addition, the predicted cue information (exteroceptive state) in the future changed, implying that the agent predicted that the food position would change in the future. Furthermore, a decrease in the interoceptive state and consequent increase in sensory uncertainty were predicted in the future. In response to the predicted danger at *t*_*c*_ = 220, the agent avoided deviating from homeostasis by resting for a while. Simultaneously, the agent planned to restart its movements while predicting increased sensory uncertainty at future time steps of approximately 400. At *t*_*c*_ = 500, the agent experienced a reduced interoceptive level and large sensory uncertainty. Future planning was generated to increase the interoceptive state and resolve deviations from homeostasis. As shown in Fig. S3, the unexpected-uncertainty-cause module adapted the posterior in the face of actual danger at *t*_*c*_ = 500, although it did not change its latent state when predicting large sensory uncertainty in the future (e.g., *t*_*c*_ = 90). This suggests that the posterior adjustment of the unexpected-uncertainty-cause module may represent the perception of uncertainty that could not be expected in advance. Eventually, by *t*_*c*_ = 900, the agent successfully survived the danger by actively obtaining food and resting again. These results show that, based on the internal predictive model, the MBH-RNN can achieve autonomous behavioral allostasis through mode switching between moving (feeding) and resting.

To analyze the mechanisms underlying behavior switching between moving and resting, we observed the behavior switching and averaged the detected network behaviors. Fig. 4 shows the changes in the posterior distributions based on the values of the 25 steps before the start of movement or rest (see Methods). The changes of posterior sigma (standard deviation) were calculated as differences from the corresponding states at the time step −25 (which can be positive or negative), while the changes of posterior mean were calculated as absolute differences from the corresponding states at the time step −25 because positive or negative sign of the change of mean states can vary depending on the different trained networks. The components of the variational free-energy (NLL and KLD terms) as well as the predicted sensory uncertainty, VFEF, and interoceptive state were also plotted. The network behaviors displayed as solid lines in Fig. 4 were analyzed using data after the end of the posterior updating process throughout the trial, meaning that the posteriors at time step *t* are the postdictive values at the tail of the past time window when the sensorimotor time step is *t*_*c*_ = *t* + *win*_*p*_ − 1 (*win*_*p*_ = 10). For reference, we plotted the immediate network behaviors recorded immediately after posterior updating at the sensorimotor time step (*t* = *t*_*c*_) as dashed lines. The postdictive posteriors displayed as solid lines at time step *t* and the immediate posteriors displayed as dashed lines at time step *t* + *win*_*p*_ − 1 are the values recorded at the same sensorimotor time step *t*_*c*_ = *t* + *win*_*p*_ − 1. In the following analyses, we focus primarily on postdictive network behavior at the tail of the past time window. This focus arises because the postdictive posteriors reflect the general network behavior across the time window, influencing the generation of forward predictions and receiving backward error signals throughout this period. Additionally, in the immediate network behaviors in Fig. 4a and 4b, the KLDs between the posteriors and priors were generally small in all modules throughout the time steps, where the priors were determined by past posteriors, including those at the tail of the past time window. Thus, we conclude that immediate network behaviors are largely controlled by postdictive posteriors.

**Fig. 4:**
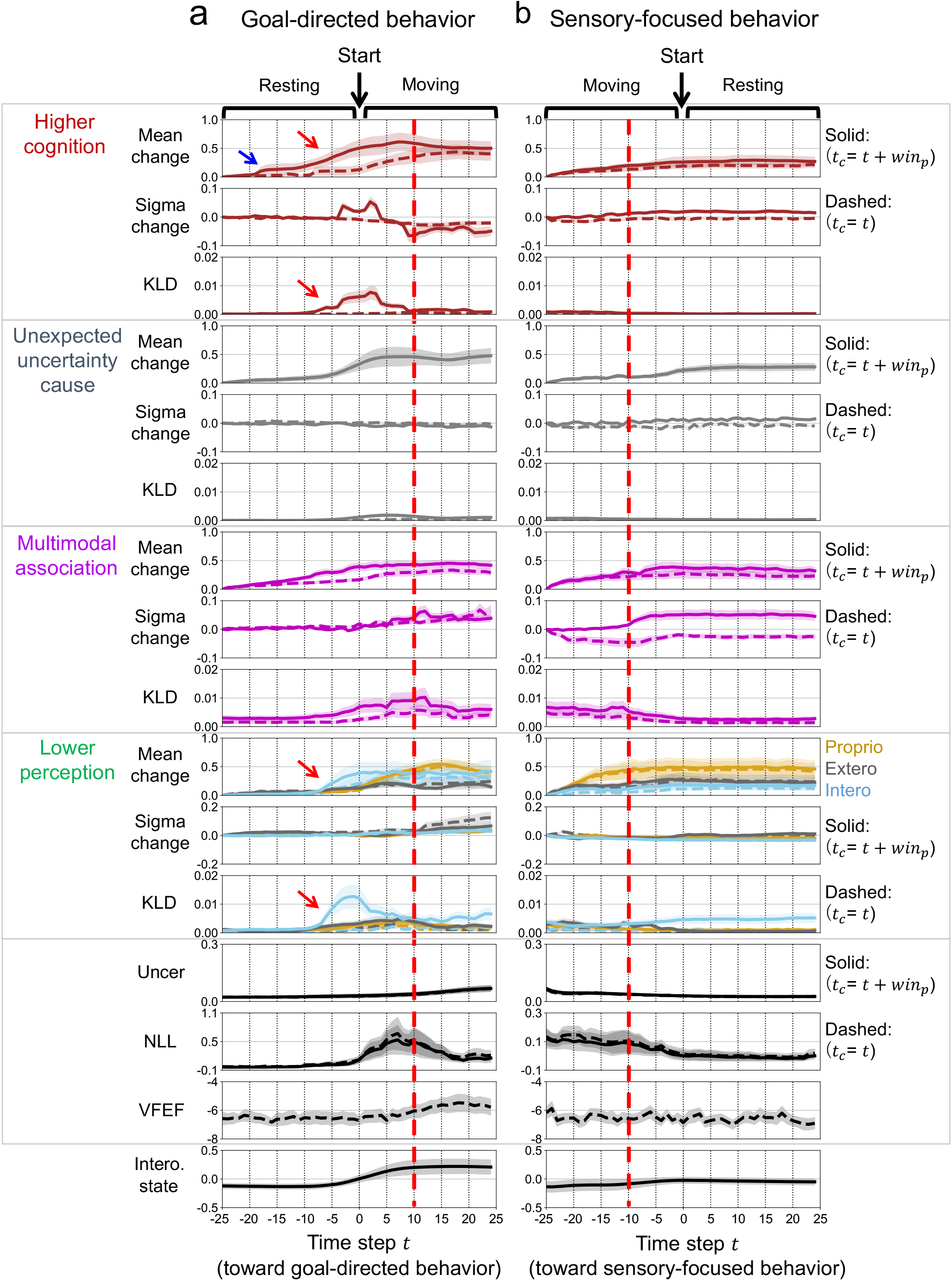
Autonomous goal-directed and sensory-focused mode switching. **(a)** Changes of posterior distributions, predicted sensory uncertainty, interoceptive state, components of variational free-energy, i.e., negative log-likelihood (NLL) and Kullback-Leibler divergence (KLD), and variational free-energy of the future (VFEF) before and after the switching from sensory-focused rest to goal-directed movement. The blue arrow indicates earlier posterior adjustments during rest in the higher-cognitive module. The red arrows indicate quick posterior changes and increased KLD signals in the higher-cognitive and interoceptive modules that triggered the goal-directed movement. The red vertical dashed line indicates increased NLL and KLDs in the multimodal-associative, proprioceptive, and exteroceptive modules but decreased KLDs in the higher-cognitive and interoceptive modules with a strong postdictive posterior in the higher-cognitive module. **(b)** The changes in the posterior distributions, predicted sensory uncertainty, interoceptive state, components of variational free-energy (NLL and KLD), and VFEF before and after switching from goal-directed movement to sensory-focused rest. The red vertical dashed line indicates the moment when the sigma of the postdictive posterior in each module started to increase. In (a) and (b), the solid linesindicate the postdictive network behaviors (*t*_*c*_ = *t* + *win*_*p*_− 1), and the dashed lines indicate the immediate network behaviors (*t*_*c*_= *t*) (*t*_*c*_: sensorimotor time step, *win*_*p*_: the length of past time window). The posterior distributions and KLDs are averaged over all dimensions of latent variables. The NLL and predicted sensory uncertainty are averaged over all sensations. The VFEF at time step *t* (*t*_*c*_ = *t*) is the value accumulated over time steps within future time window *t*_*c*_ + 1: *t*_*c*_ + *win*_*f*_ (*win*_*f*_: the length of future time window). These values are also averaged over all sequences extracted from timeseries of 10 test trials in each network. Line shadings represent the standard error over 10 different trained networks. Uncer: predicted sensory uncertainty. Proprio: proprioceptive module. Extero: exteroceptive module. Intero: interoceptive module.

Fig. 4a shows the behavior switching from resting to moving. First, before the start of movement, the postdictive posterior (solid line) in the higher-cognitive module was slightly adjusted around *t* = − 18 as indicated by the blue arrow (see also the slight adjustment of the immediate posterior displayed as a dashed line in that module at around the corresponding sensorimotor time step *t*_*c*_ = −9). During rest, the levels of NLL (sensory prediction errors) were very small because of the limited interaction with the environment, suggesting that earlier posterior adjustments may result from goal-directed errors for food intake (via the VFEF). Then, right before the start of moving, quick adjustments of postdictive posteriors in the higher-cognitive and lower interoceptive modules were observed from around *t* = −9, with increases of KLDs in the corresponding modules. These postdictive posterior adjustments may be ascribed to the gradual increase in VFEF after the corresponding sensorimotor time step *t*_*c*_ = 0. The start of the movement for food intake was reflected in the change in the posterior of the proprioceptive module and an increased interoceptive state after *t* = 0. During movement, the postdictive posteriors in the higher-cognitive and interoceptive modules were maintained, and the sigma of the postdictive posterior in the higher-cognitive module decreased. The time step of the posterior sigma in the higher-cognitive module hit the bottom (around *t* = 10), coinciding with the moment when the VFEF reached its peak (around the corresponding sensorimotor step *t*_*c*_ = 19). This suggests that large goal-directed errors form a strong (postdictive) belief with a low estimated sigma. Importantly, the movement and resultant interaction with the environment after *t* = 0 led to increases in the NLL and KLDs in the multimodal-associative, proprioceptive, and exteroceptive modules, whereas the KLDs in the higher-cognitive and interoceptive modules decreased. In other words, the MBH-RNN modulates which components of the VFE should be minimized to realize future homeostatic and interoceptive goals. Mode switching from resting to moving may involve the dynamic regulation of the strength (precision) of hierarchical latent states, in which the higher-cognitive and interoceptive modules maintain strong postdictive posteriors, ignoring sensory prediction errors during movement. As such, the emergence of strong beliefs or “intention” enabled movement generation at the expense of large sensory prediction errors.

In contrast, Fig. 4b shows the behavior switching from moving to resting. In this case, the interoceptive state was regulated to an appropriate level before the agent began to rest. At the moment of the start of resting, the sigma of the postdictive posterior in each module was generally increased after around *t* = −10 (the corresponding sensorimotor step is *t*_*c*_ = −1), as highlighted by the vertical red line. The start of rest was reflected in the invariant posterior position of the proprioceptive module after *t* = 0. In addition, the rest greatly decreased the NLL, which may be ascribed to a reduction in the interaction with the environment. These observations suggest that the weak postdictive posteriors with high estimated sigma served as sensory-focused “recognition” of internal and external states that were adapted towards minimizing sensory prediction errors, leading to the rest. In summary, Fig. 4a and 4b illustrate how autonomous switching between goal-directed and sensory-focused modes emerges through recognition-intention transitions in Bayesian beliefs.

#### Predictive interoceptive regulation

We investigated whether the Bayesian allostasis model could explain the increase in food intake in preparation for food shortage periods, as observed in biological cognitive agents^1,4,5^. For clarity, we compared the proposed allostasis model with a setpoint model (control condition) that had an explicit homeostatic setpoint (target value) of the future interoceptive state set to 0.0. In the setpoint model, the predicted sensory entropy term of the VFEF is replaced by a NLL for the interoceptive sensation, given a fixed setpoint or the prediction error between the predicted future interoceptive sensation and the setpoint. Equivalent interoceptive prediction errors given a homeostatic setpoint have historically been explored to explain goal-directed homeostatic processing within various contexts, including cybernetics, active inference (with the concept of expected free energy), and homeostatic reinforcement learning^49–52^. Note that we used the same trained MBH-RNNs for both the allostasis and setpoint models, meaning that the difference was only in the autonomous survival process after learning.

We first evaluated the difference in the survival time steps between the allostasis and setpoint models (Fig. 5a). A paired t-test revealed that the allostasis model survived significantly longer than the setpoint model did (*t*(9) = 3.38, *p* = 0.00*T*2). To analyze the general characteristics of interoceptive regulation in the survival task, the cycle of food nutrition was divided into five periods: the food recovery period (I), the food gradually decreasing periods (II-IV), and the food shortage period (V) (Fig. 5b). Fig. 5c shows the difference between the two models in interoceptive regulation by plotting the changes in the interoceptive state during the food cycle based on the values in period I. The allostasis model increased the interoceptive level until Period III, and the interoceptive level in Period V was similar to that in Period I. Simultaneously, the number of food acquisitions peaked in Period III, following an increased acquisition of cue information in Period II (Fig. 5d). Which indicate that the allostasis model exhibits predictive interoceptive regulation by capturing the food nutrition cycle and exploring food positions through information gain. However, in the setpoint model, the interoceptive level, as well as the acquisition of food and cue information, monotonically decreased during the food gradually decreasing periods II-IV, and the interoceptive level in Period V was smaller than that in Period I (Fig. 5c and 5e). Furthermore, the allostasis model showed a smaller amount of energy expenditure throughout the food nutrition cycle than the setpoint model did, which may have increased the effectiveness of energy intake (Fig. 5f). Differences between the allostasis and setpoint models in the regulation of energy intake and expenditure may result in significant differences in survival performance.

**Fig. 5:**
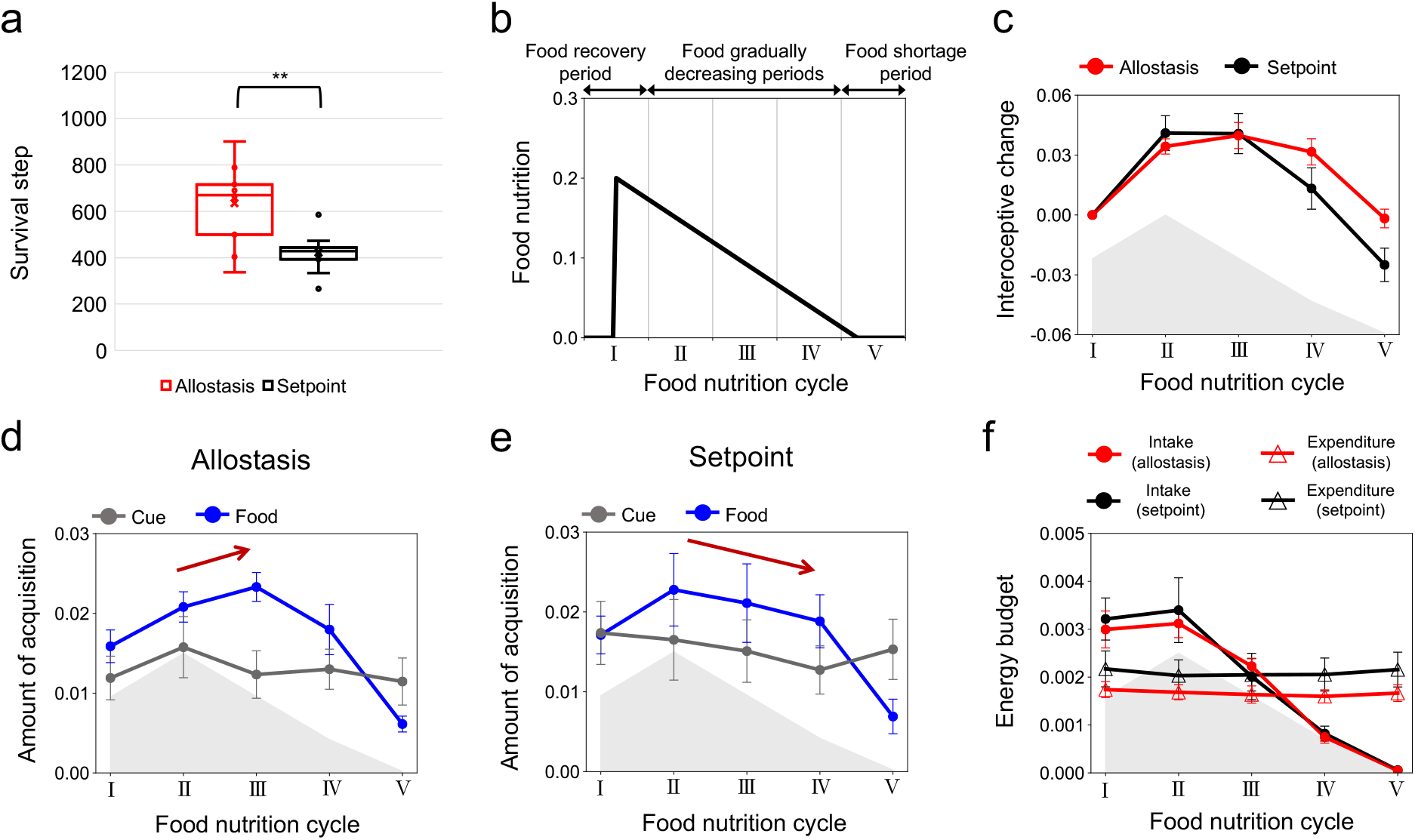
Predictive interoceptive regulation. **(a)** Box-and-whiskers plots showing the difference in autonomous survival time step between the allostasis model and the setpoint model (control condition). Each plot is an average of the 10 test trials by each of 10 trained networks (center line, median; cross, mean; box limits, upper and lower quartiles; whiskers, × 1.5 interquartile range). ** *p* < 0.01. **(b)** the food nutrition cycle is equally divided into five periods: from food recovery period (I) to food gradually decreasing periods (?-?), and food shortage period (V). **(c)** Change of interoceptive state based on the value in the period I in each model. **(d**,**e)** Changes in the amounts of food and cue acquisitions per time step in the allostasis model and the setpoint model. **(f)** Changes of energy intake and energy expenditure per time step in the allostasis model and the setpoint model. In (c,d,e,f), the gray shadow represents the shape of the cycle of averaged food nutrition for reference, and error bars represent the standard error.

## 3. Discussion

We propose that an autonomous cognitive agent does not just minimize sensory prediction errors but also minimizes predicted future sensory entropy. We demonstrated that this Bayesian allostasis model can be derived by extending the variational Bayesian framework to future predictive processing. We validated the Bayesian allostasis model by simulating a structurally and temporally hierarchical MBH-RNN that exhibited self-organizing behavior, switching between goal-directed movement (feeding) and sensory-focused rest, based on the internal predictive model. The continuous mode switching resulted in predictive interoceptive regulation in a dynamic environment, which may explain behavioral allostasis in biological cognitive agents^1,4,5^. Our Bayesian neural network framework opens new avenues for exploring brain information processing to anchor continual behavioral adaptation.

Goal-directed and sensory-focused mode switching were involved in switching the roles of Bayesian beliefs in VFE minimization. During the goal-directed movement, strong postdictive posteriors were maintained and served as “intention.” However, in the sensory-focused rest, weak postdictive posteriors were adapted towards minimizing sensory prediction errors, serving as “recognition” of internal and external states. It may be reasonable to precisely predict the future risks of restarting feeding. These results suggest that mode switching correspond to meta-level modulations in which components of the variational free energy are minimized by the agent. This self-organization of mode switching may explain how “recognition” changes into “intention” and vice versa in a Bayesian brain.

In particular, switching from sensory-focused rest to goal-directed movement may demonstrate how cognitive agents actively interact with the external environment, even if it is unpredictable or surprising. At the moment of the mode switching, generations of large KLD signals (due to goal-directed signals for food intake) changed the higher-level postdictive posteriors into strong beliefs (“intention”), which triggered movements for feeding. The observation of adjustments of the higher-level postdictive posteriors during rest may be associated with biological observations of the readiness potential in advance of conscious intention to action^53,54^, which might be associated with different internal experiences of the agent in terms of the experienced level of goal-directed signals. This may have computational and phenomenological implications for the emergence of intentions.

Our Bayesian allostasis model suggests that the homeostatic behaviors of cognitive agents may primarily be guided by the aspiration of a certain computation concerning second-order prediction (low predicted future sensory entropy) rather than maintaining physical states at the setpoints. The previously proposed setpoint-driven models for homeostasis and allostasis (explained as adjustments of setpoints)^49–52^ can be considered a specific case of the goal-directed mode in our model. In addition, our model explains how strong beliefs regarded as “setpoint” or “intention” can emerge in the hierarchical neural network. From another perspective, our model integrates the concept of reward in reinforcement learning into the Bayesian brain hypothesis by considering interoceptive predictive processing in the face of increased physiological entropy owing to irreversible physical processes within the body.

Autonomous mode switching remains a largely unexplored phenomenon in computational modeling studies of the brain. However, it may be a key for studying “intention” and “self” in the computational machine. Biological and theoretical studies have suggested that different kinds of sensory suppression processes, such as sensory attenuation and gating, may be associated with self-nonself distinction and a sense of agency^23,46^. Sensory attenuation refers to fewer perceptual and neural responses to self-generated sensations than to externally generated sensations, whereas sensory gating refers to fewer responses to sensations during movement than during rest^55^. Sensory attenuation can be explained by self-organized mode switching in the process of VFE minimization, depending on the inferred causality of sensations (self or nonself)^20,46^.

Specifically, in the mechanism of sensory attenuation through VFE minimization, constantly strong Bayesian beliefs at a lower perceptual level were observed during the self-generated context. This led to a higher weight of the corresponding KLD terms at the lower perceptual level than those at the higher network levels, resulting in reduced prediction-error-induced responses at the lower perceptual level. However, the current model study showed that emergence of strong beliefs at the interoceptive and higher-cognitive modules enabled movement generation by ignoring sensory prediction errors, where the corresponding KLD terms (associated with the strong beliefs) of VFE were weighted higher than NLL term as well as KLD terms in other modules. Ignoring sensory prediction errors due to strong beliefs in the goal-directed mode may explain sensory gating during movement. These observations provide mechanistic implications for the difference between sensory attenuation and gating, although both can be explained as autonomous mode switching in VFE minimization. The self-organization of autonomous mode switching may pave the way for studying the computational structures underlying autonomy in terms of perception and behavior.

Neuroimaging studies suggest that allostatic-interoceptive processing involves a large-scale brain network that is composed of core regions of the salience and default mode networks, such as the insula, anterior cingulate cortex, prefrontal cortex, amygdala, thalamus, and hypothalamus^56^. Based on the lines of analysis of our trained model (Fig. 2–4 and S3), we speculate about a correspondence between the functional modules of our MBH-RNN and the allostatic-interoceptive system in a biological brain. Regarding the sensorimotor modules of the MBH-RNN, the interoceptive module represented modality-specific interoceptive information, whereas the multimodal-associative module integrated interoceptive information with exteroceptive and proprioceptive information. This hierarchical interoceptive neural representation may be analogous to the hierarchical functional organization of the insular cortex^37^. In contrast, regarding homeostatic modules, the higher-cognitive module of the MBH-RNN integrated multimodal information with sensory-uncertainty information and controlled goal-directed behavior. This functionality may be consistent with biological knowledge about cognitive control regions. The characteristics of the unexpected-uncertainty-cause module, perception of uncertainty and unchanged latent state, might be associated with brain regions representing a range of uncertainty information, such as the amygdala and basal forebrain. In summary, the self-organizing functional network of the MBH-RNN may provide insights into the functional architecture of the allostatic-interoceptive network^57^ and related regions in the brain.

As a limitation, we focused only on the cognitive aspect of allostasis, which requires behavioral interactions with the external environment. Further discussions may be required for interoceptive controls via reflex arcs such as cardiovascular properties (e.g., blood pressure), which seem to be involved with predefined reactions in the central nervous system based on “hard-wired” setpoints^58^. Additionally, future studies should explore the scalability of our model in a physical or simulated robot with high-dimensional sensors, as well as validate the *in silico* neural network model with a biological body. Furthermore, the application of our model to explore the computational mechanisms underlying altered interoceptive processing in psychiatric disorders (e.g., depression, eating disorders, and autism spectrum disorder)^59–61^ and to investigate possible interventions for preventing or improving them will be a potential direction for future studies.

## 4. Methods

### Experimental setup

In this section, we explain the settings used to create the sensory data. The radius of the food and cue fields was set to 0.2, which was used to judge the acquisition. The relationship between the real standard deviation of sensory noise *σ*_*x,t*_ and the interoceptive state *x*_*t,intero*_ (the effect of deviations from homeostasis) is defined as the summation of sigmoid functions (the graph can be seen in Fig. 1a).

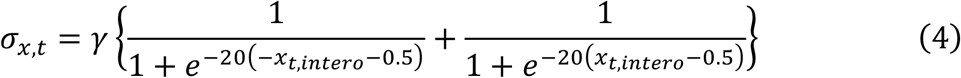

Here, 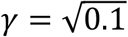 determines the maximum of *σ*_*x,t*_. In biological agents, the relationship between *σ*_*x,t*_ and *x*_*t,intero*_ may be determined by the bodily characteristics of the agent, such as muscles and metabolism, which are regulated through bodily changes (allodynamics). Although the relationship was fixed in our experiment, we confirmed that allostatic interoceptive regulation was observed even in a smaller *γ* setting 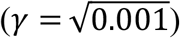, showing the robustness of our allostasis model (Fig. S4).

The energy expenditure (decrease in the interoceptive state) EE_*t*_ owing to movement is calculated at each time step based on the moving distance MD_*t*_ (difference between the current agent’s position and the previous position) as EE_*t*_ = *k*(0.1 + MD_*t*_). Here, *k* = 0.013 controls the level of expenditure and is heuristically determined such that the average interoceptive state in the learning data is nearly 0.

### Neural network model

In this section, we describe the mathematical details of top-down prediction generation and bottom-up parameter updates using the MBH-RNN, which is based on PV-RNN^46^ (Fig. S1).

Prediction generation was performed in a top-down manner using a network hierarchy. The internal state *h*_*t,i*_ and output *d*_*t,i*_ of the *i*th deterministic variable (recurrent unit) in each network module at time step *t* (*t* ≥ 1) is calculated as

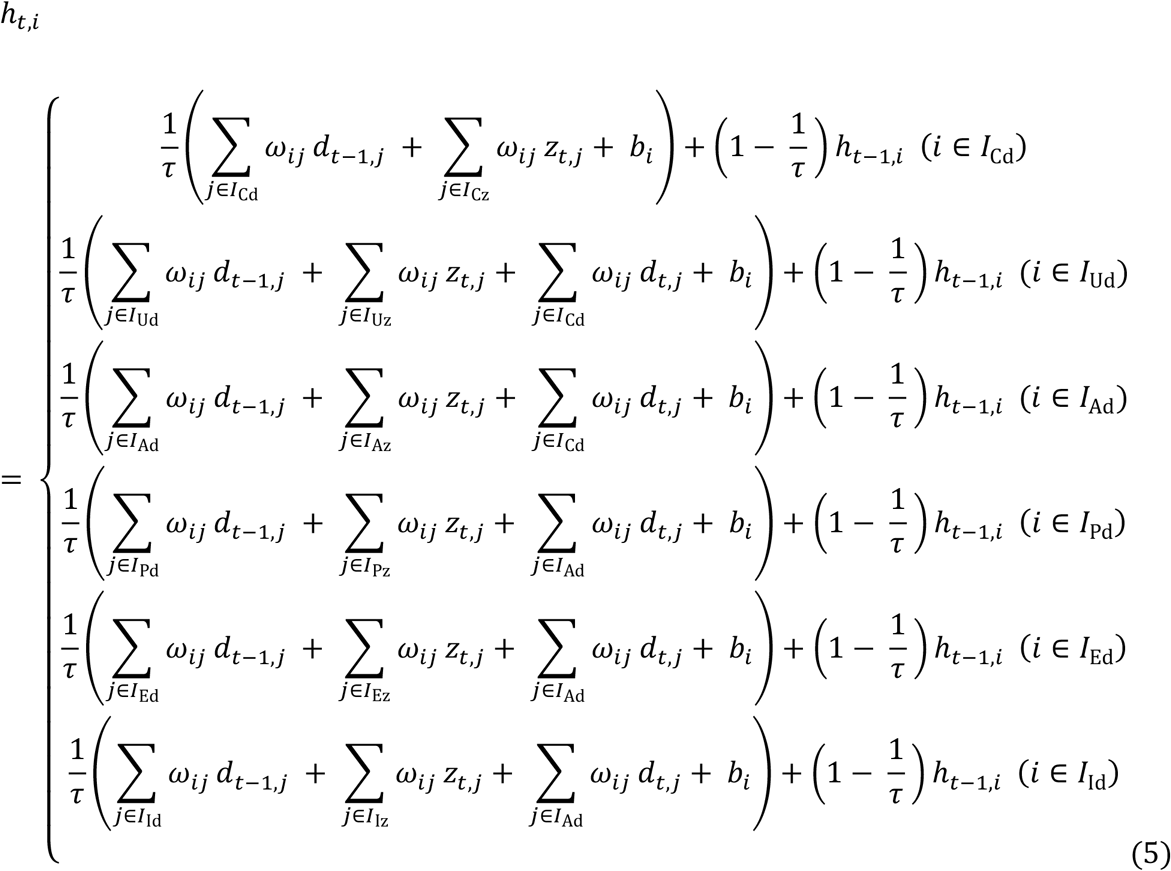

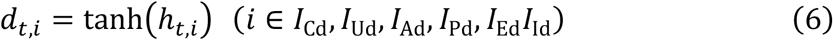

where *I*_Cd_, *I*_Ud_, *I*_Ad_, *I*_Pd_, *I*_Ed_, and *I*_Id_, are the index sets of deterministic variables in the higher-cognitive module, unexpected-uncertainty-cause module, multimodal-associative module, proprioceptive module, exteroceptive module, and interoceptive module, respectively. *I*_Cz_, *I*_Uz_, *I*_Az_, *I*_Pz_, *I*_Ez_, and *I*_Iz_ are the index sets of probabilistic latent states in the higher-cognitive module, unexpected-uncertainty-cause module, multimodal-associative module, proprioceptive module, exteroceptive module, and interoceptive module, respectively. *ω*_*ij*_ is the weight of the synaptic connection from the *j*th neuron to the *i*th neuron; *z*_*t,j*_ is the output of *j*th latent states at time step *t*; τ is the time constant of the deterministic variable; and *b*_*i*_ is the bias of the *i*th deterministic variable. A deterministic variable with a small time constant τ has a tendency to change its activity rapidly, while that with a large time constant has a tendency to change its activity slowly^24^. We set the initial internal states of the deterministic variables *h*_0,*i*_ (*i* ∈ *I*_Cd_, *I*_Ud_, *I*_Ad_, *I*_Pd_, *I*_Ed_, *I*_Id_) to 0 (*d*_*T,i*_ is also 0).

The probabilistic latent state ***z***_*t*_ in each module is assumed to follow a multivariate Gaussian distribution with a diagonal covariance matrix, meaning *z*_*t,i*_ and *z*_*t,j*_ are independent (*i, j* ∈ *I*_*C*z_, *I*_Uz_, *I*_Az_, *I*_Pz_, *I*_Ez_, *I*_Iz_ ∧ *i* ≠ *j*). Here, the mean *μ*_*p,t,i*_ and sigma (standard deviation) *σ*_*p,t,i*_ of the prior distribution *p*(*z*_*t,i*_) in all modules are calculated as below:

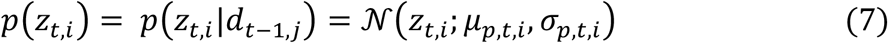

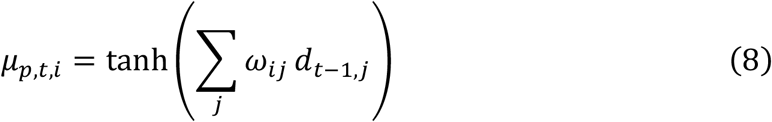

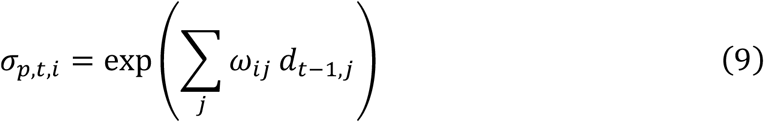

Here, prior distributions in each module are calculated from the previous deterministic states of the same module, meaning (*i* ∈ *I*_Cz_ ∧ *j* ∈ *I*_Cd_) ∨ (*i* ∈ *I*_Uz_ ∧ *j* ∈ *I*_Ud_) ∨ (*i* ∈ *I*_Az_ ∧ *j* ∈ *I*_Ad_) ∨ (*i* ∈ *I*_Pz_ ∧ *j* ∈ *I*_Pd_) ∨ (*i* ∈ *I*_Ez_ ∧ *j* ∈ *I*_Ed_) ∨ (*i* ∈ *I*_Iz_ ∧ *j* ∈ *I*_Id_).

The posterior distribution in each module is calculated as,

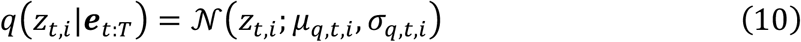

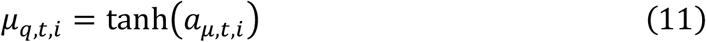

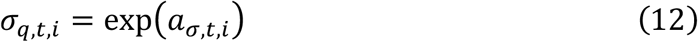

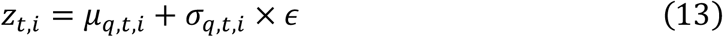

Here, *T* represents the last time step of the time window. ***a***_*t*_ is the adaptive internal state of the latent variables representing the posterior distributions. The adaptive variables ***a***_*t*_ are determined by error signals ***e***_*t*:*T*_ propagated using the BPTT algorithm. The adaptive variables ***a***_*t*_ are initialized by the corresponding initial internal states of the latent variables that represent the prior distributions. In equation (13), by sampling *ϵ* from *𝒩*(0,1), the latent state *z*_*t,i*_ is obtained.

Finally, mean predictions about proprioceptive, exteroceptive, and interoceptive sensations were individually generated from the proprioceptive, exteroceptive, and interoceptive modules, respectively, whereas sigma predictions for each sensation were generated from the unexpected-uncertainty-cause module.

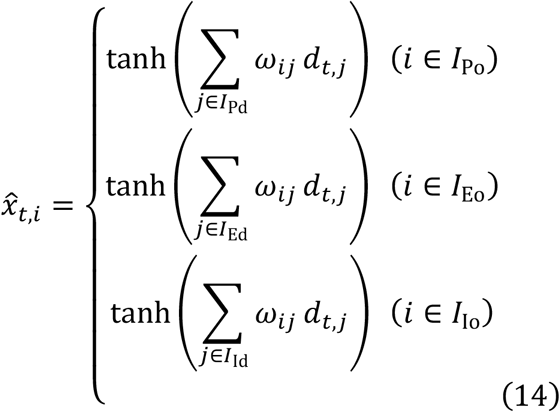

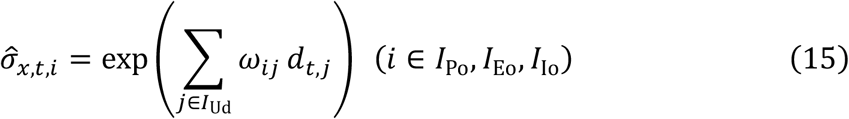

Here, *I*_Po_, *I*_Eo_, *I*_Io_ are the index sets of the output units for the proprioceptive, exteroceptive, and interoceptive predictions, respectively.

The cost function of the MBH-RNN is derived based on variational Bayes or the free-energy principle (FEP)^8^. Under the FEP, minimizing the surprisal (or the negative log model evidence) for sensations ***x*** over time is the ultimate goal of self-organizing systems. In the learning of the MBH-RNN, the surprisal over all time steps can be written as:

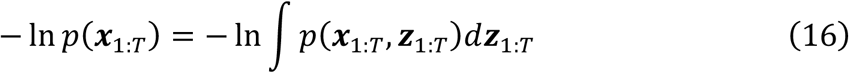

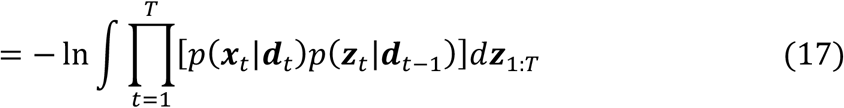

Here, surprisal cannot be directly evaluated by the agent because it needs to know all hidden states ***z*** of the environment that cause sensations, as described on the right side of the equation (16). Here, FEP introduces a tractable quantity, VFE, that bounds the surprisal, and minimization of the surprisal is replaced by minimization of the VFE. The bound of the surprisal in the MBH-RNN can be derived by utilizing Jensen’s inequality for a concave function *f*: *f*(*E*[*x*]) ≥ *E*[*f*(*x*)]. Then, Equation (17) can be modified as follows by introducing a dummy distribution for ***z***_1_, *q*(***z***_1_).

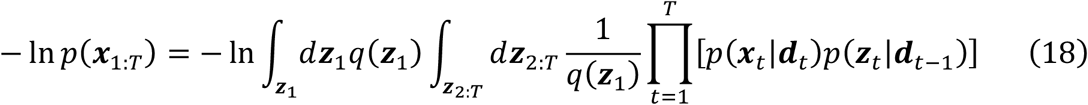

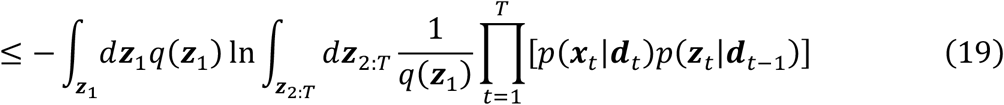

In Equation (19), Jensen’s inequality is used as a logarithmic function. The same procedure is performed for *t* = 2: *T*.

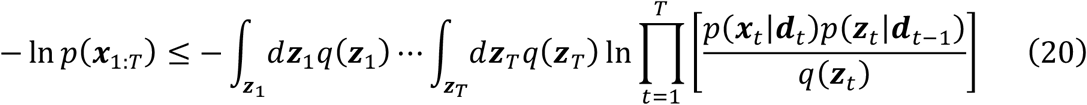

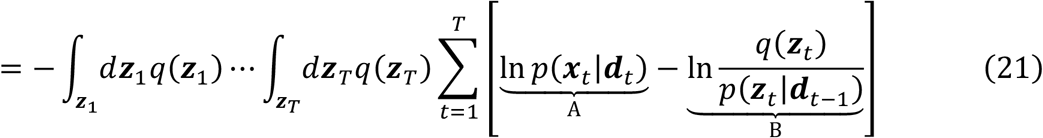

The first term in expression (21) is the expected NLL under *q* given that ***d***_*t*_ depends on ***z***_1:*t*_.

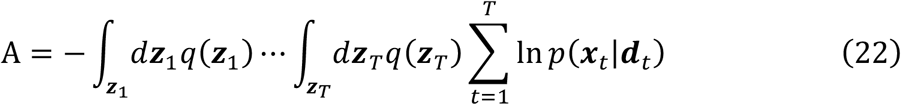

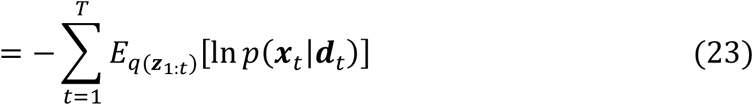

In addition, the second term can be deformed into forms of KLD between *q*(***z***_*t*_) and *p*(***z***_*t*_|***d***_*t*−1_) while ensuring that ***d***_*T*_ is independent of ***z***_1:*T*_.

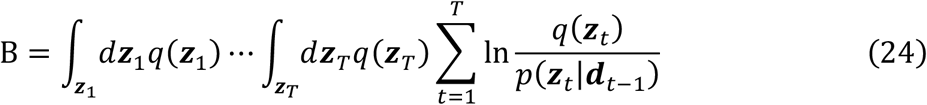

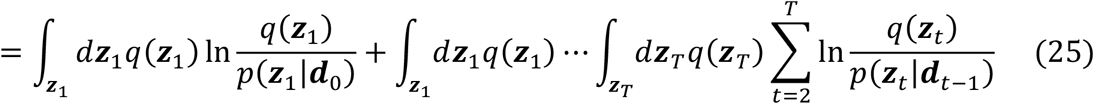

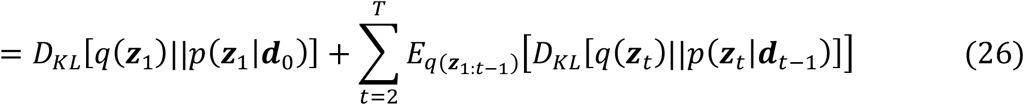

A dummy distribution *q*(***z***_*t*_) can be replaced by the posterior distribution determined by the back-propagated prediction error *q*(***z***_*t*_|***e***_*t*:*T*_). Thus, using equations (23) and (26), the bound of the surprisal can be written as,

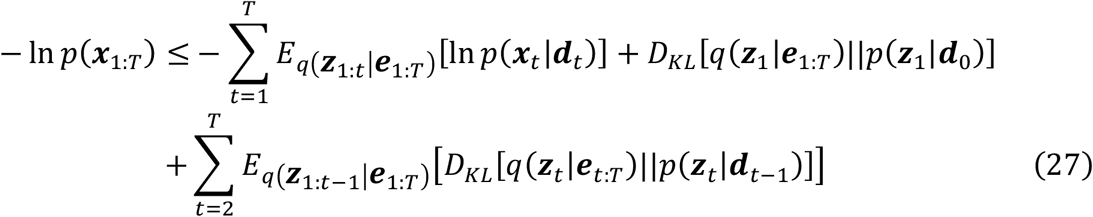

In the experiment, we resolved the calculation of the expectations of the NLL and KLD under *q* by considering a single sampling to reduce computational costs.

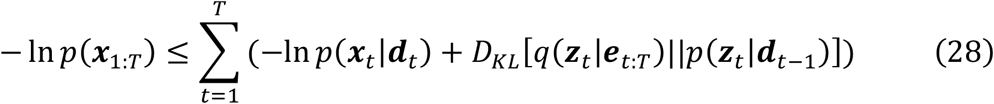

Eventually, the cost function (the bound of the surprisal) in the learning *F*_*learning*_ is the accumulation of variational free-energy VFE_t_ over time steps.

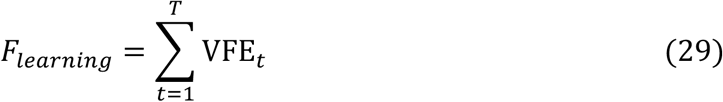

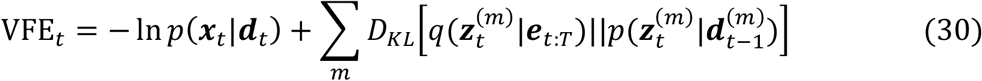

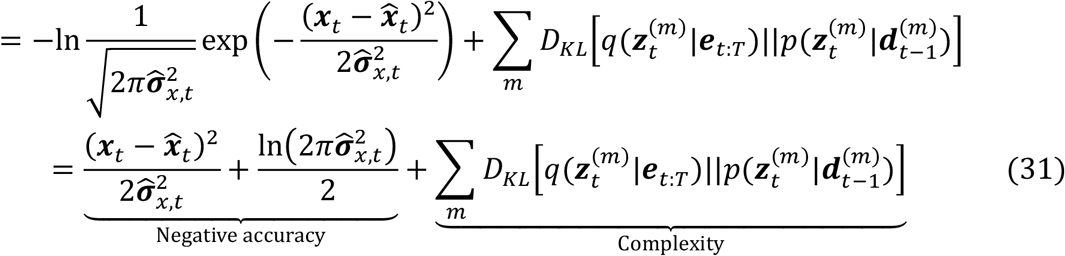

where *m* denotes a network module. The first term in expression (31) (negative accuracy term) is the NLL. For simplicity, we assume that each sensory state follows a Gaussian distribution. In the experiment, we divided the negative accuracy term by the dimensions of the proprioceptive, exteroceptive, and interoceptive sensations. However, the second term (complexity term) is the KLD between the posterior and prior distributions of the latent states. Assuming that the prior and posterior distributions follow a multivariate Gaussian distribution with a diagonal covariance matrix, as described above, the KLD is computed analytically as^45^,

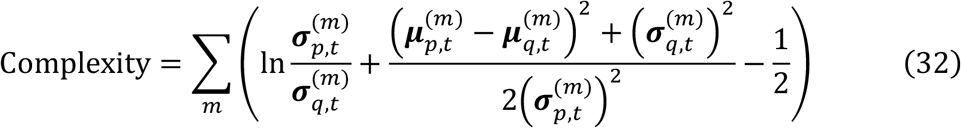

In the experiment, we divided the complexity term by the dimensions of the latent variables for each module.

In the learning phase, synaptic weights ***ω*** and adaptive variables ***a***_1:*T*_ are updated to minimize the VFE_*t*_ over all time steps (*T* = 10,000). We used gradient descent with a rectified Adam optimizer^62^ for parameter updates, where the partial derivative of the variational free energy with respect to each parameter was calculated using the BPTT algorithm.

By contrast, in the autonomous survival task, the MBH-RNN performed posterior updates within time steps from *t*_*c*_ − *win*_*p*_ + 1 to *t*_*c*_ + *win*_*f*_ at each sensorimotor time step *t*_*c*_ (Fig. S2). The cost function in the past time window *F*_*past*_ is the accumulated VFE of the past based on the observed sensations 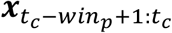.

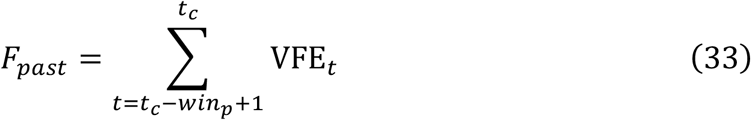

In contrast, VFEF is derived by considering a bound on the expected negative log model evidence (or the expected surprisal) under the distribution of predicted future sensory trajectories.

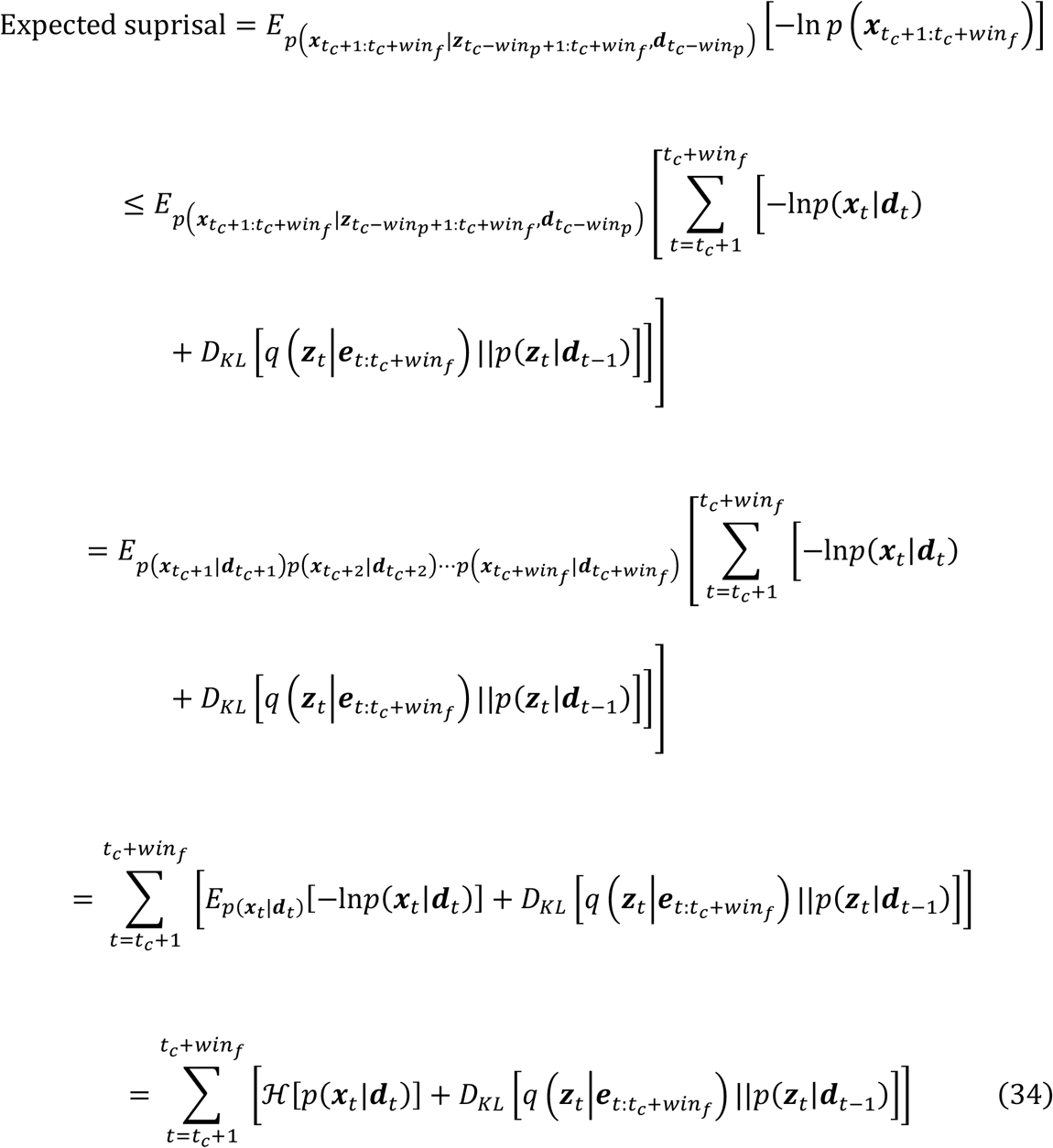

Finally, the cost function in the future time window, *F*_*future*_ is derived by considering the KLD term in each module.

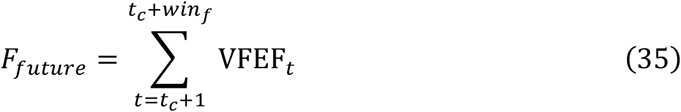

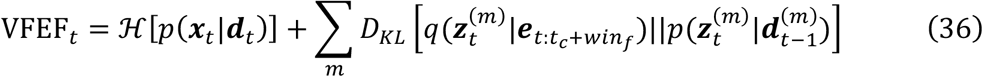

Here, the first term is the conditional (Shannon) entropy of the sensations, and is calculated as follows by assuming a Gaussian distribution:

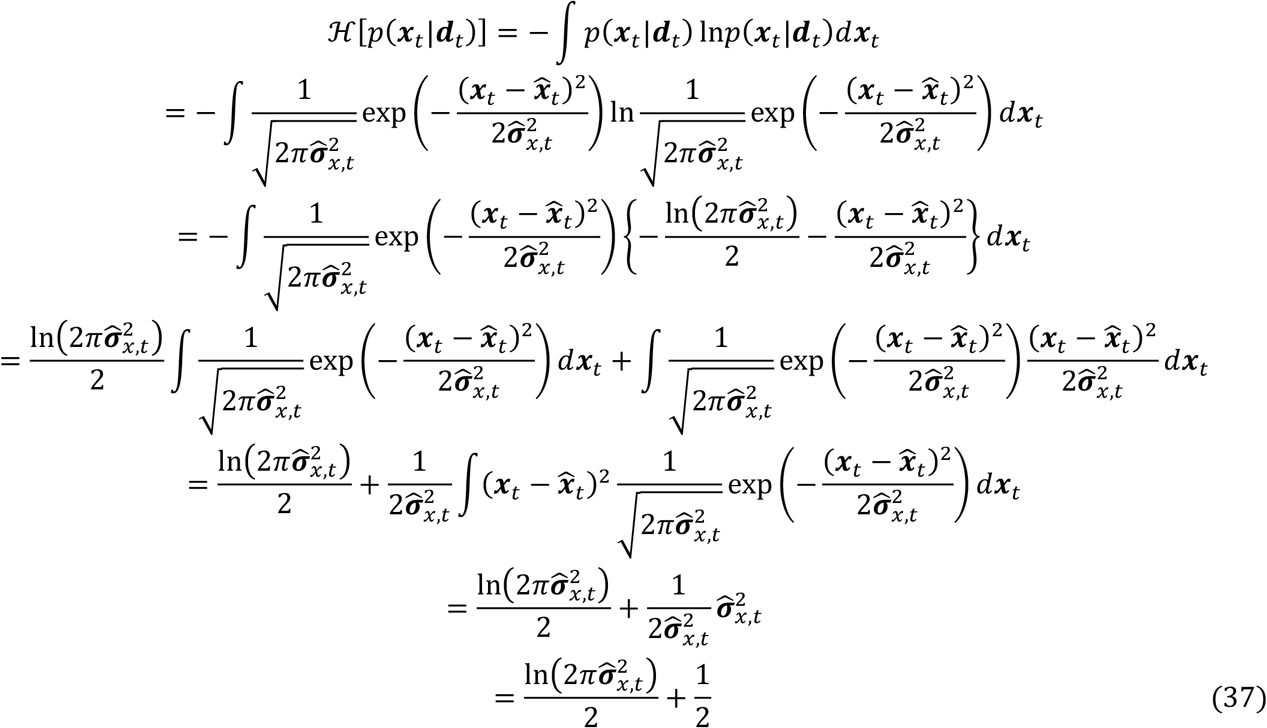

Thus, the VFEF is described below.

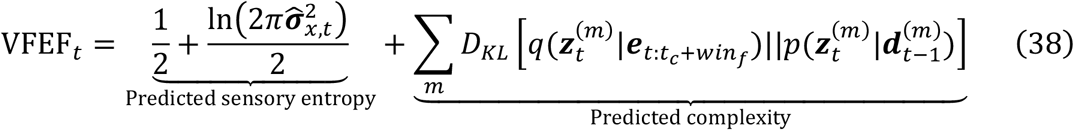

This derivation of VFEF naturally describes the minimization of the predicted future sensory entropy (uncertainty) or the meta-goal of Bayesian allostasis.

The MBH-RNN performs iterations of prediction generations and posterior updates within time steps from *t*_*c*_ − *win*_*p*_ + 1 to *t*_*c*_ + *win*_*f*_ to minimize the accumulated free-energy from the past to the future *F*_*allostasis*_ through BPTT.

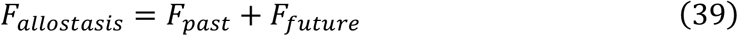

Furthermore, we prepared a setpoint model as the control condition in which the predicted sensory entropy term of the VFEF was replaced by a NLL (predicted negative accuracy term) given the explicit interoceptive target state (homeostatic setpoint) 0.0, which is equivalent to the prior preference for future outcomes within the concept of active inference applied to explain goal-directed interoceptive control^50^.

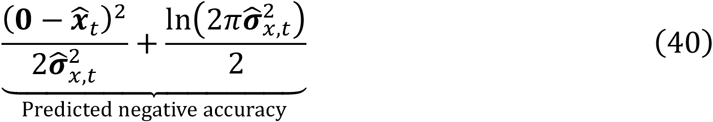

### Hyper-parameter setting

The dimensions of the latent variables in the proprioceptive, exteroceptive, and interoceptive modules were *N*_Pz_ = 2, *N*_Ez_ = 2, *N*_Iz_ = 1, respectively, corresponding to the dimensions of the sensations in each modality. The dimension of latent variables in the unexpected-uncertainty-cause module was *N*_Uz_ = 1 as the minimal setting, while that in the higher-cognitive module was *N*_Cz_ = 3 as the dimension necessary for associating the information of each of the three modalities with sensory uncertainty. The dimension of the latent variables in the multimodal-associative module was *N*_Az_ = 8, which was determined by a preliminary experiment in which we increased the dimension of the latent variable in the multimodal-associative module individually and searched for the minimal setting for successful reconstruction of the learning data in all trained networks with different initial synaptic weights. For simplicity, the numbers of deterministic recurrent variables in the proprioceptive, interoceptive, unexpected-uncertainty-cause, and higher-cognitive modules were *N*_pd_ = *N*_Id_ = *N*_Ud_ = *N*_Cd_ = 10 for simplicity, while those in the exteroceptive and multimodal-associative modules were *N*_Ed_ = *N*_Ad_ = 20 required for reconstructing the learning data. We incorporated a temporal hierarchy (multiple timescale properties) into the MBH-RNN. Specifically, for each of the lower perceptual modules, we set the time constant as τ_Pd_ = τ_Ed_ = τ_Id_ = 2 or 4 (for each of half the number of deterministic recurrent variables) as the simple multiple timescales setting. Then, the time constants in the multimodal-associative module and unexpected-uncertainty-cause module were set as τ_Ad_ = τ_Ud_ = 4 or 8 (for each of the half-deterministic recurrent variables), while those in the higher-cognitive module were set as τ_Cd_ = 8 or 16 (for each of half deterministic recurrent variables). As such, the higher-level modules have a larger time constant (slower neural dynamics) than the lower-level modules have, thus implementing a temporal hierarchy. The synaptic weights were initialized with random values based on the Xavier method^63^. The biases of the deterministic recurrent variables were initialized with and fixed to random values following a Gaussian distribution *𝒩*(0,10), based on a previous study^17^. For quantitative evaluation, we trained 10 networks with different initial synaptic weights and biases for each hyper-parameter setting. In the learning phase, parameters including synaptic weights and adaptive (posterior) variables were updated *itr*_*learning*_ = 100,000 times with the rectified Adam optimizer (learning rate *lr*_*learning*_= 0.01, *β*_1_ = 0.9, *β*_2_ = 0.999). In the autonomous survival task test, adaptive variables were updated *itr*_*inference*_ = 200 times at each sensorimotor time step *t*_*c*_ for a time window from the past time window of length *win*_*p*_ = 10 (*win*_*p*_ = *t*_*c*_ *if t*_*c*_ < 10) to the future time window of length *win*_*f*_ = 200 with *lr*_*inference*_ = 0.1. We chose the optimal parameter settings in the survival task from the combinations of *lr*_*inference*_ = {0.01, 0.05, 0.1, 0.2, 0.5, 1.0} × *itr*_*inference*_ = {100, 200, 500} × *win*_*p*_ = {10, 20, 50} × *win*_*f*_ = {50, 100, 200} by evaluating the autonomous survival time steps.

### Analysis of latent representation

An analysis of the developed latent representation via the ablation of latent variables was conducted, as shown below (Fig. 2c). We regard the network predictions (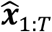and 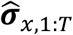) at the end of learning as the target of network predictions. First, for calculating the baseline variability of the network predictions, we made the trained normal network generating 10 different timeseries (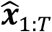 and 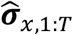) with different sets of random sampling of latent states. The differences between the target timeseries and the 10 re-generated timeseries were calculated, and the maximum value of the differences for each of the proprioceptive, exteroceptive, interoceptive, and uncertainty predictions was named the baseline difference (BD). Next, we removed one latent variable and made the lesioned network generating 10 different timeseries (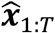 and 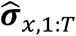) with different random sampling. The differences between the target timeseries and the 10 timeseries generated by the lesioned network were calculated, and the average of the differences for each of the proprioceptive, exteroceptive, interoceptive, and uncertainty predictions was named the ablation effect (AE). If the AE for a certain modality (proprioceptive, exteroceptive, interoceptive, or uncertainty predictions) was larger than the corresponding BD was, we considered that the ablated latent variable represented the affected modality information. For example, if the ablation of a latent variable of the multimodal-associative module affects proprioceptive and interoceptive sensory predictions, the ablated latent variable is considered to represent both proprioceptive and interoceptive information and is a bimodal latent variable. We repeated the evaluation for all latent variables in all network modules for all 10 trained networks. Finally, we categorized the types of latent variables based on the information represented in each network module and calculated the probability for each type of latent variable.

### Analysis of the dynamics of posteriors and free-energy components

In Fig. 4a, we extracted the behavior switching from resting to moving, as shown below: First, we calculated the distance moved at each time step using the time series obtained from each test trial. For all time steps *t* (>25), the averages of the moving distances over the time steps from *t* − 25 to *t* − 1 and from *t* to *t* + 24 were regarded as MD_before_ and MD_after_, respectively. If MD_before_ was lower than one-third of MD_after_ and the moving distance at each time step from *t* − 25 to *t* − 1 were all smaller than 0.5, we considered that the switching from resting to moving started at time step *t*. In contrast, in Fig. 4b, if MD_after_ was lower than one-third of MD_before_ and the moving distance at each time step from *t* to *t* + 24 were all smaller than 0.5, we considered that the switching from moving to resting started at time step *t*. Finally, in each trained network, we averaged the extracted network behaviors for all behavior switchings observed in 10 test trials.

### Statistical analysis

We used paired t-tests for statistical analyses to compare the proposed allostasis model and the setpoint model (control condition), which were tested using the same trained MBH-RNNs. All statistical tests were two-tailed, and the significance level was set at *p* < 0.05. In this study, an unprecedented computational simulation was conducted. Thus, it was difficult to estimate the effect size, and no statistical methods were used to pre-determine the sample size. Considering the high reproducibility of the computational simulation, we set the minimum sample size that seemed statistically testable (10 samples). Indeed, even for the 10 samples, the paired t-test reported clear differences in the analyses. Therefore, we conclude that a larger sample size would not have significantly influenced our main results. Data analyses were conducted using the R software (version 3.3.2).

## Acknowledgements

HI acknowledges the support from a Grant-in-Aid from the Japan Society for the Promotion of Science Research Fellows (Nos. JP22J01708, JP22KJ3167). YY acknowledges support from the Japan Science and Technology Agency Moonshot R&D (No. JPMJMS2031), and grants from the Japan Science and Technology Agency Core Research for Evolutional Science and Technology (No. JPMJCR16E2, JPMJCR21P4). TO acknowledges support from the Japan Science and Technology Agency Moonshot R&D (No. JPMJMS2031). The funders had no role in the writing of the manuscript or in the decision to submit the manuscript for publication.

## Data Availability

All data is available in the manuscript and the supplementary information.

## Author contributions

Conceptualization: H.I.; methodology: H.I.; software: H.I.; investigation: H.I.; resources: H.I. and Y.Y.; visualization: H.I., J.T., and Y.Y.; funding acquisition: H.I., T.O., and Y.Y.; project administration: H.I., Y.Y.; supervision: J.T., T.O., and Y.Y.; writing—original draft: H.I.; writing—review and editing: H.I., J.T., T.O, and Y.Y.

