## Supplementary figures for "Future shapes present: autonomous goal-directed and sensory-focused mode switching in a Bayesian allostatic network model"

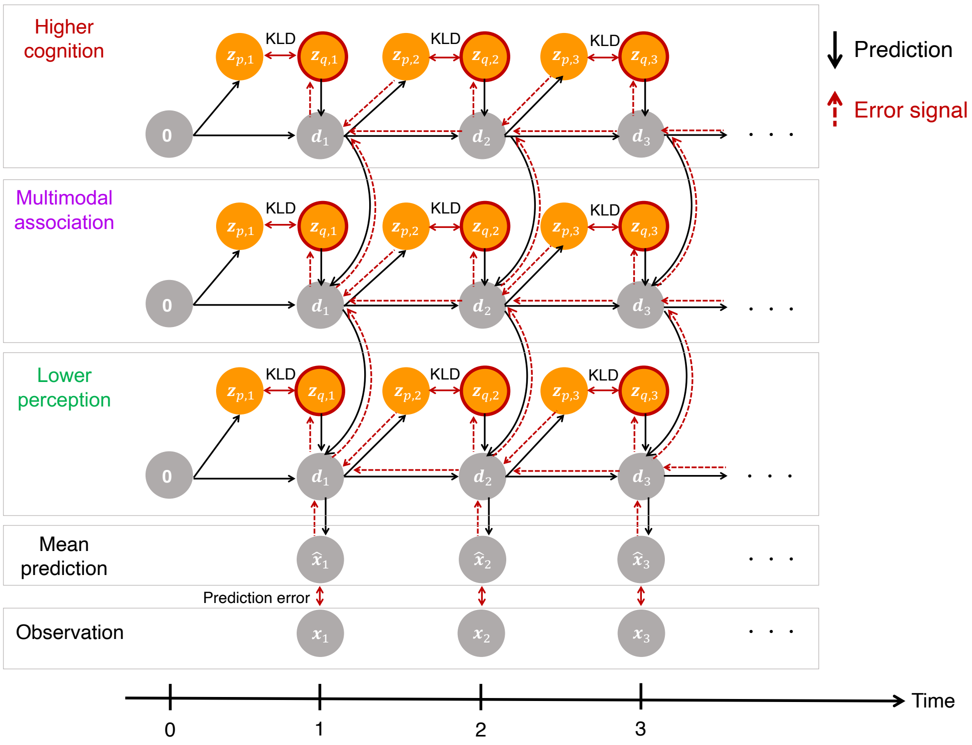


**Fig. S1: Temporal processing of the MBH-RNN**

Related to Fig. 2a. In the learning process, the MBH-RNN updates the posterior variables $\boldsymbol{a}_{1:T}$ in all modules and time-invariant synaptic weights $\boldsymbol{\omega}$ via minimization of variational free-energy over the time length of the learning data $T$. The initial deterministic states in all modules are set to 0. For simplicity, the distributed structures of the lower-perceptual modules as well as the unexpected -uncertainty-cause module and sensory-uncertainty prediction are omitted. KLD: Kullback-Leibler divergence between posterior and prior latent variables.

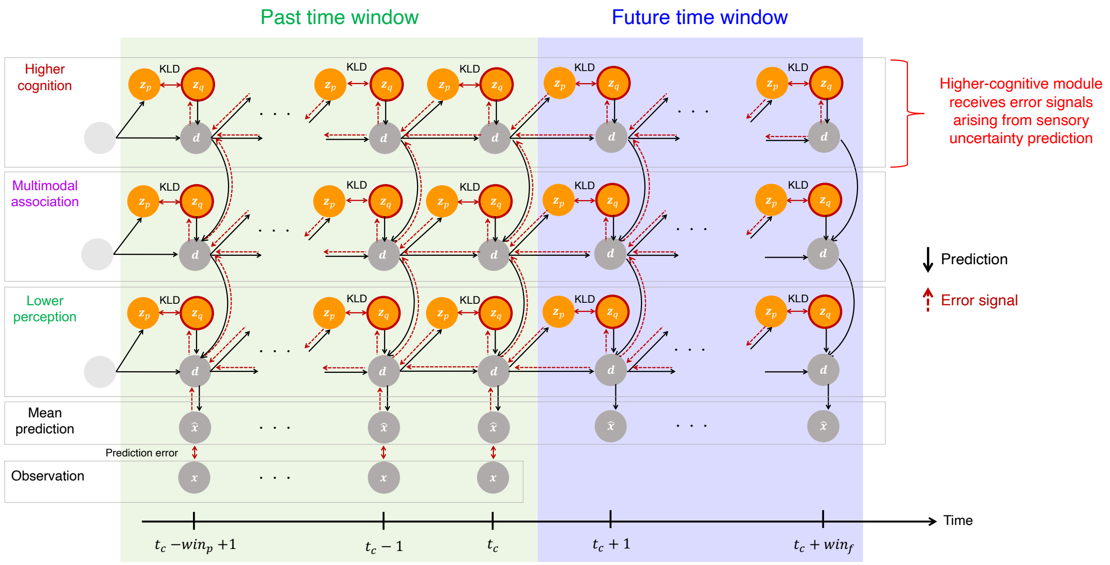


**Fig. S2: Online inference in the survival test**

Related to Fig. 3a. At the current sensorimotor time step $t_c$, the MBH-RNN updates the posterior variables $\boldsymbol{a}_{t_c - win_p + 1 : t_c + win_f}$ in all modules via minimization of the variational free-energy from the past to the future. The synaptic weights are fixed. For simplicity, the distributed structure of the lower-perceptual modules as well as the unexpected-uncertainty-cause module and sensory-uncertainty prediction are omitted. KLD: Kullback-Leibler divergence between the posterior and the prior of the latent variable.

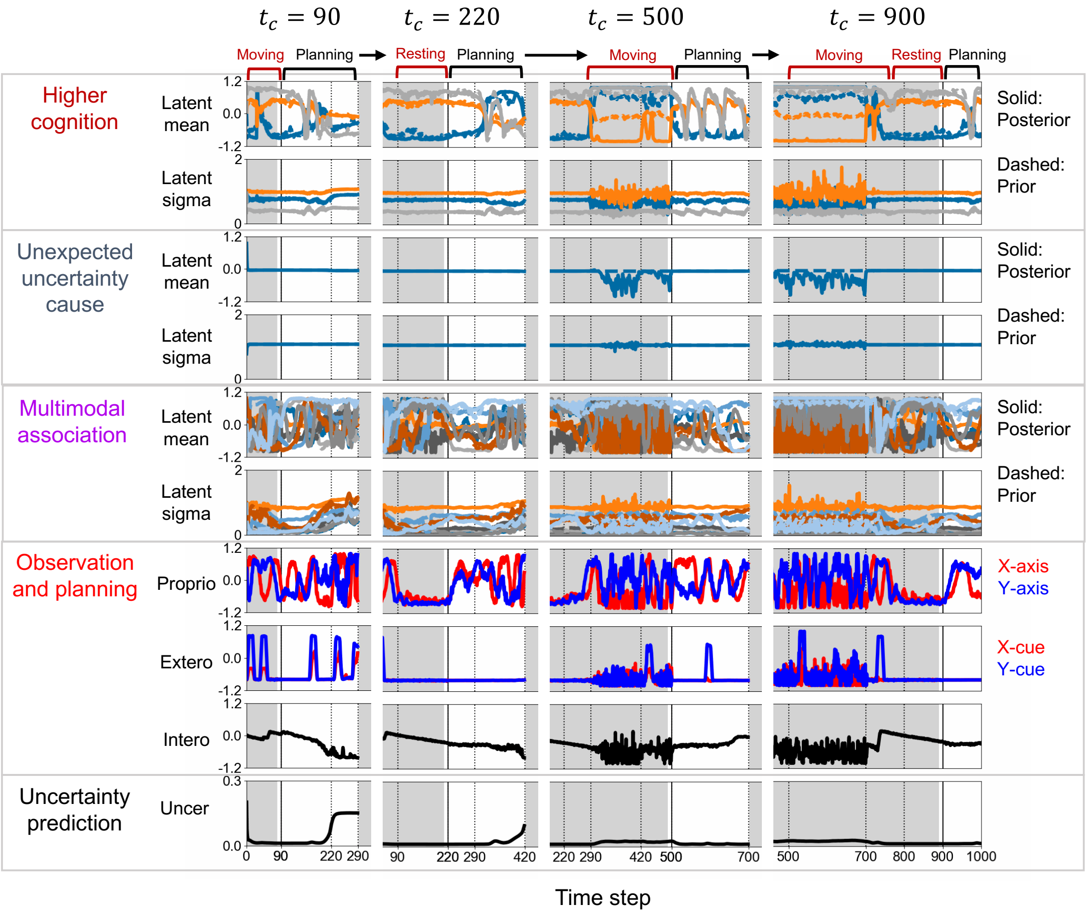


**Fig. S3: Supplementary result for latent dynamics during autonomous survival task**

Related to Fig. 3b. Prior and posterior distributions in the higher-cognitive module, unexpected-uncertainty-cause module, and multimodal-associative module are added to Fig. 3b. An important point is that the latent states of the higher-cognitive module changed according to the homeostatic context rather than the short-term modality-specific context. In addition, the latent state of the unexpected-uncertainty-cause module did not change when predicting large sensory uncertainty in the future (e.g., $t_c = 90$) but responded to actual increases in sensory uncertainty or deviations from homeostasis (e.g., $t_c = 500$ ). This shows that the unexpected-uncertainty-cause module represented the cause of sensory uncertainty that could not be expected in advance.

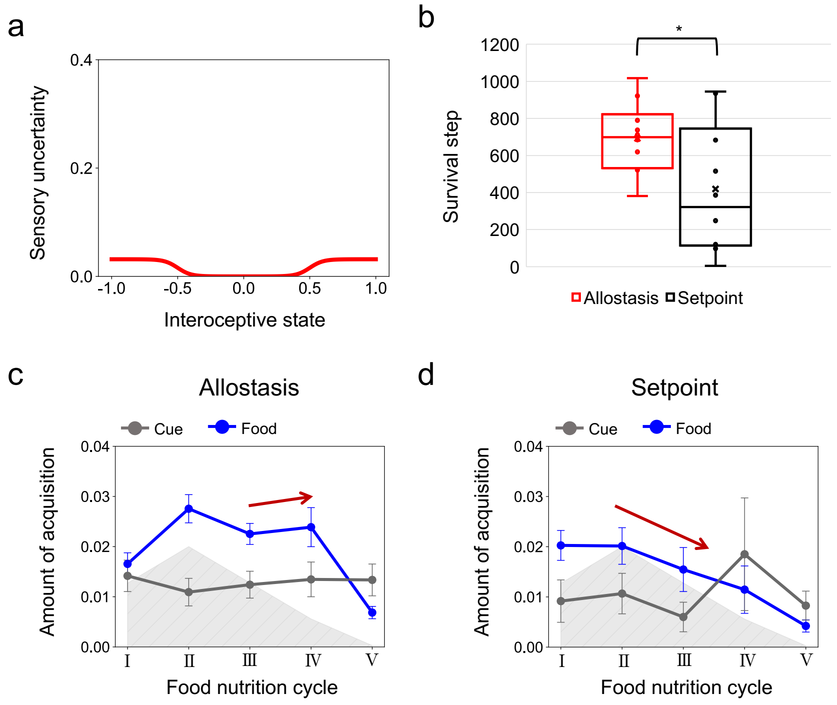


**Fig. S4: Supplementary results for different settings of the relationship between sensory uncertainty and the interoceptive state**

**(a)** The maximum of sensory uncertainty (standard deviation) is set to $\gamma = \sqrt{0.001}$, a smaller value than the setting investigated in the main text ($\gamma = \sqrt{0.1}$). **(b)** Box-and-whiskers plots showing the difference in autonomous survival time step between the proposed allostasis model and the setpoint model (control condition). A paired t-test reported that the allostasis model survived significantly longer than the setpoint model did ($t(9) = 2.36, p = 0.043$). Each plot is an average of the 10 test trials by each of 10 trained networks (center line, median; cross, mean;

box limits, upper and lower quartiles; whiskers, $\times$ 1.5 interquartile range). $* p < 0.05$. **(c,d)** Changes in the amounts of food and cue acquisitions per time step in the allostasis model and the setpoint model. The allostasis model increased the food acquisitions in the period Ⅳ before the food shortage period Ⅴ while the setpoint model exhibited a monotonical reduction of food acquisitions during the food gradually decreasing periods Ⅱ-Ⅳ. These results show the robustness of our Bayesian allostasis model. The gray shadow represents the shape of the cycle of averaged food nutrition for reference, and error bars represent the standard error.

### Caption for supplementary videos

**Video S1.** Related to Fig. 3b and S3. Agent behavior and dynamics of latent states during autonomous survival tasks.
